# The Intrinsic Hierarchy of Self – Converging Topography and Dynamics

**DOI:** 10.1101/2022.06.23.497287

**Authors:** Yasir Çatal, Mehmet Akif Günay, Chunbo Li, Jijun Wang, Huiru Cui, Wei Li, Georg Northoff

## Abstract

The brain can be characterized by an intrinsic hierarchy in its topography which, as recently shown for the uni-transmodal distinction of core and periphery, converges with its dynamics. Does such intrinsic hierarchical organization in both topography and dynamic also apply to the brain’s inner core itself and its higher-order cognitive functions like self? Applying multiple fMRI data sets, we show how the recently established three-layer topography of self (internal, external, mental) is already present during the resting state and carried over to task states including both task-specific and -unspecific effects. Moreover, the topographic hierarchy converges with corresponding dynamic changes (measured by power-law exponent, autocorrelation window, median frequency, sample entropy, complexity) during both rest and task states. Finally, analogous to the topographic hierarchy, we also demonstrate hierarchy among the different dynamic measures themselves according to background and foreground. Finally, we show task-specific- and un-specific effects in the hierarchies of both dynamics and topography. Together, we demonstrate the existence of an intrinsic topographic hierarchy of self and its convergence with dynamics.

## 1 Introduction

The brain’s intrinsic organization shows a complex topographic hierarchy. One topographic key feature is the distinction of uni- and transmodal regions. Unimodal regions at the brain’s periphery include the sensory regions which can be characterized by a hierarchical organization among themselves (like primary, secondary, etc. sensory regions) [37, 62, 72]. In contrast, transmodal regions like the default-mode network constitute the core [42, 48, 67, 71, 74]. What remains unclear is whether, analogous to the periphery, there is hierarchical organization among the regions within the inner core itself.

The transmodal core is associated with higher-order cognition like episodic simulation [26, 28, 49], mind wandering which are closely related to our self [40, 68, 78, 79] featured by self-referential processing [13, 20, 21, 43, 54].

The self is a multifaceted higher-order cognitive feature of our mind which includes interoceptive, bodily, and mental features [13, 23, 36, 54]. Each of these facets has been associated with different regions in the brain like insula [70], temporoparietal junction (TPJ) [23, 32], and cortical midline structure (CMS) [10, 13, 17, 54]. This raises the question of their relationship and ultimately the topographic organization of these various regions. Is there an intrinsic hierarchical organization of self within the brain’s topography of its inner core? Addressing this yet unresolved question is the goal of our paper.

A recent meta-analysis of task-related studies on self-specificity yielded three distinct layers of self: the internal (interoceptive) self is related to the insula, dorsal anterior cingulate cortex, and thalamus; the external (exteroceptive-proprioceptive) self is mediated by the premotor cortex, TPJ and the mental self recruits with anterior and posterior CMS [59]. Importantly, these different regions are embedded and thus nested within each other with the lower hierarchy regions (like internal self regions e.g. insula and anterior cingulate) being recapitulated in the next higher layer of the external self which recruits additional regions (like TPJ) and so forth. Together, this amounts to a multi-layered topographic hierarchy of self. This leaves open whether hierarchical organization of self is truly intrinsic. In that case, one would expect the three-layered hierarchical organization of self to be present already in the spontaneous activity itself as measured during resting state, that is, during the absence of task-evoked activity related to self-specific stimuli or other self-referential contents.

Complementing its topographic organization, various EEG and fMRI studies demonstrate specific dynamic features characterizing the self. These include stronger power in slow frequencies measured by the power-law exponent (PLE) as well as longer intrinsic neural timescales operationalized by the autocorrelation window (ACW) [41, 51, 56, 81]. Complementing the dynamics of brain activity, one can also measure its degree of information processing by for instance sample entropy [7, 30, 65, 69, 72] and Lempel-Ziv Complexity (LZC) [3, 8, 11, 74]. How the dynamic of self (and its information processing) are related to and possibly converge with the intrinsic three-layered topographic hierarchy of self remains yet unclear, though.

The goal of our paper is to investigate whether the self exhibits an intrinsic topographic hierarchy within the brain’s inner core itself and how that converges with its dynamics. This is stipulated by recent studies showing that the intrinsic core-periphery topography converges with dynamic resulting in a slow core (with long timescales) and faster periphery (with short timescales) [55, 60, 67, 74]. Based on these studies we hypothesized that the three-layered topographic hierarchy of a the self is intrinsic e.g. holds during both rest and task states, and is met by a more or less corresponding hierarchy in its dynamics. Specifically, we hypothesized the longest autocorrelation (ACW) and strongest slow frequency power (PLE) to hold in the mental layer of self (as related to the CMS as the topographic core of the brain). While the internal layer of self, being more distant from the core may show shorter autocorrelation (ACW) and less power in slow frequencies (PLE).

The first specific aim consists in calculating the dynamic and information-based measures (ACW, PLE, MF, LZC, SE) in the regions of the three-layer topography of self during the resting state. Based on the assumed intrinsic nature of the topographic hierarchy and its convergence with dynamics, we hypothesized different degrees in these measures within the regions of the three layers of self. More generally, following the data on core-periphery [67, 74], we assumed that the three-layer topography of self converge with corresponding differences in the measures that index dynamics (ACW, PLE, MF) and information processing (SE, LZC).

The second specific aim consists in probing the convergence of dynamics and topography of self during a variety of task states in order to reveal task-specific and unspecific effects in dynamics. For that purpose, we calculated our various dynamic measures in the regions of the three-layer topography during 7 different task states from two datasets. We hypothesized that the three-layer hierarchical organization of self is carried over from rest to task states thus further underlining its intrinsic nature – this accounts for largely task-unspecific effects. While task-specific effects can be observed in case the task itself targets one particular layer of self, e.g., internal, external, or mental.

The third specific aim consists in characterizing and operationalizing the hierarchical organization of self by itself, that is, its degree of hierarchy in both its topographic (relation among the three layers) and dynamic (relationship among the different measures) terms. For that purpose, we calculated a gradient among the three layers of self and hypothesized that that gradient changes during a self-specific task (whereas that may not be the case if one applies tasks unrelated to the self). Moreover, applying the graph-theoretic measures, we investigated the relationship among the five dynamic measures. We hypothesized that the dynamic measures like PLE, MF, and ACW are more stable during both rest and task states, operating in the background while information-related measures like SE and LZC are more flexible showing changes during task states thus taking on the role as foreground measures [82].

Applying multiple fMRI rest and task data sets, we show an intrinsic topography of a higher-order cognitive function like self within the brain’s core as well as its convergence with dynamics during both rest and task states; the topographic differences of internal, external, and mental self during the resting state was accompanied by corresponding differences in dynamics, probed with PLE, ACW, MF; and information processing measured by LZC and SE. This was also apparent during the task state suggesting carry-over of the topographic hierarchy from rest to the task that further supports the intrinsic nature of the brain’s topography of self. When subtracting task from rest, we were able to distinguish task-specific and -unspecific effects in both topography and dynamics. Finally, applying graph-theoretic measures, we demonstrate that dynamic measures (ACW, PLE, MF) constitute a hub of high centrality and closeness during both rest and task states. In contrast, information processing measures (SE, LZC) were more flexible showing higher changes in their centrality during task states. Together, our findings show, for the first time, an intrinsic hierarchical topography for a higher-order cognitive function like the self within the brain’s inner core and its convergence with a corresponging dynamic hierarchy.

## 2 Methods

The codes to replicate preprocessing, calculation of measures and doing the network analysis with simulated signals can be found at https://github.com/duodenum96/Self-Topography-Dynamics. All statistical tests were corrected for multiple comparisons using Bonferroni-Holmes method. asterisks denote significance; *: *p <* 0.05, **: *p <* 0.01, ***: *p <* 0.001. Effect size estimates for all of the Kruskal-Wallis and Wilcoxon tests were calculated in R [77] using rstatix [64] package and can be found in supplementary material. Tables in supplementary materials were generated with help of the packages gt [75], tidystats [80], broom [73] and tibble [76].:

### 2.1 Data Acquisition and Task Designs

Two different fMRI datasets were used in this study. The first dataset (will be referred to as the Shanghai dataset from now on), [53] included 50 age and sex-matched subjects. fMRI images were acquired on a 3.0-T Siemens MAGNETOM Verio syngo MR B17 scanner equipped with a 12-channel head coil (Siemens, Erlangen, Germany). T1-weighted images were as follows: repetition time (TR), 1,900 ms; echo time (TE), 2.46 ms; inversion time (TI),900 ms; flip angle (FA), 9°; field of view (FOV), 256 × 256 mm; matrix, 256 × 256; slice thickness, 1 mm (no gap); and 192 sagittal slices. Four echo-planar imaging (EPI) scans were obtained for task fMRI and one EPI scan was obtained for rest fMRI. The sequence parameters were as follows: TR, 2,000 ms; TE, 32 ms; FA, 70°; FOV, 240 × 240 mm; matrix, 64 × 64; slice thickness, 5 mm; 30 interleaved transverse slices; voxel size, 3.8 × 3.8 × 5 mm. Event-related task design [9, 15, 22, 39] included 3 conditions: rest, interoceptive stimulus (listening to one’s own heartbeat) and exteroceptive stimulus (listening and counting auditory tones), 9-13 seconds each. Each of the 4 runs lasted 9.6 minutes. The measures calculated from different runs were averaged to get one measure per subject in the task condition. For a general linear model analysis as well as detailed information regarding subjects, we refer the readers to [53].

The second fMRI dataset (slice thickness = 4 mm, 34 slices, TR = 2 s, TE = 30 ms, flip angle = 90?,matrix 64 × 64, FOV = 192 mm,oblique slice orientation, will be referred as UCLA dataset from now on) was downloaded from open access UCLA Consortium for Neuropsychiatric Phenomics LA5c study [44, 45]. Only raw data of 130 healthy controls were selected for analysis. Details about sociodemographic characteristics of the sample, detailed MRI device information, anatomical scan parameters and information on resting state and 6 tasks can be found on [44, 45]. Briefly, these tasks consist of an eyes open resting state (REST); balloon analog risk task (Bart), in which subjects were asked for pumping experimental balloons that either resulted in an explosion or points and control balloons which neither exploded nor rewarded; paired associate memory encoding (Pamenc) and retrieval (Pamret), in which subjects were asked for memorization and rate their confidence in recalling the memorized color; spatial working memory task (Scap) in which subjects tried to remember places of pseudo-randomly positioned circles; stop-signal task (Stopsignal), in which subjects tried to not respond to the visual cue when they hear an auditory signal; and task-switching task (Taskswitch), in which participants were asked for responses to either shapes or colors of the stimuli.

### 2.2 Preprocessing

Preprocessing steps were implemented in Analysis of Functional Neuroimages software (AFNI; [5]). The procedure is following: 1) discarding the first two frames in UCLA and 5 frames in each Shanghai dataset of each fMRI run to ensure proper magnetization; 2) slice-timing correction; 3) despiking; 4) spatial alignment of fMRI data to time frame with minimum estimated motion; 5) spatial alignment of fMRI data to skullstripped anatomical data that was aligned to MNI152 stereotactic space; 6) resampling to 3 × 3 × 3 mm isometric voxels; 7) scaling each voxel time series to have a mean of 100 to reflect percent signal change; 8) temporal band-pass filtering (0.01 Hz ¡ f ¡ 0.2 Hz) to reduce low-frequency drift and high-frequency respiratory/cardiac noise, while at the same time undesired components were removed through regression of linear and nonlinear drift: head motion and its temporal derivative, binarized Euclidean norm time series of motion above the threshold of 2 mm, mean time series from the white matter (WM) and cerebrospinal fluid (CSF) to control for non-neural noise. The WM and CSF masks were eroded by one voxel to minimize partial voluming with gray matter [19]. Censored time points were linearly interpolated to keep temporal continuity. Global signal regression was not performed to avoid introducing non-existent correlations and losing correlations due to global signal, which is shown to have physiological relevance [27, 52, 61, 63, 66].

After preprocessing, the quality control files generated by afni proc.py were visually inspected and the specific runs from the subjects with excessive motion and artifacts were discarded. This left us with the following number of subjects for each task: for the Shanghai: 39 rest and 43 task scans; for the UCLA, 108 for Rest and Scap, 107 for Bart, Stopsignal and Taskswitch, 75 for Pamenc and Pamret.

### 2.3 ROI Selection

The ROIs were selected from [59]. The three layers of self were operationalized using ROIs from figure 7 in the paper (table 1). The BOLD signals inside spheres with a diameter of 8 mm were averaged to get one signal per ROI and the measures calculated for these ROIs were averaged to get one measure per layer using AFNI’s 3dcalc and 3dmaskave.

**Table 1:** ROI selection

| ROI | x | y | z |
| --- | --- | --- | --- |
| Internal |  |  |  |
| LIns1 | -40 | -2 | 2 |
| LIns2 | -36 | 24 | 4 |
| RIns | 34 | 14 | 12 |
| External |  |  |  |
| MPFC | -6 | 60 | 22 |
| TPJ | -48 | -38 | 36 |
| PMC | 50 | 8 | 26 |
| Mental |  |  |  |
| PACC | -6 | 48 | 0 |
| PCC | -4 | -54 | 28 |

ROIs were selected from [59]. The BOLD signal inside spheres with 8 mm diameter were averaged to get one BOLD signal per ROI. After the calculation of measures, they were averaged across layers to get one measure per layer. Moreover, we overlaid our ROIs and the transmodal networks (as determined in [55]) of the atlas from [50]. This showed us that our self layers are inside the core (transmodal) instead of periphery (unimodal) of the brain’s organization of intrinsic neural timescales. This visualization can be seen in supplementary figure 1.

### 2.4 Calculation of measures

All the measures were calculated on MATLAB software (version R2020a). We calculated power-law exponent (PLE), median frequency (MF), autocorrelation window (ACW-0), sample entropy (SE) and Lempel-Ziv complexity (LZC) to probe the dynamics of self layers.

#### 2.4.1 Power-Law Exponent and Median Frequency

Power spectral density (PSD) of frequencies between 0.01 and 0.2 Hz were calculated using the periodogram method [4, 12]. The periodograms were smoothed with a Hamming shaped sum-cosine function based on AFNI’s 3dTsmooth with -hamming option. PLE was defined as the slope of the linear regression line that fits the power versus frequency representation in logarithmic scale [18, 29] to get the exponent *β* in 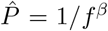 where *P* is power, *f* is frequency and *β* Is the power-law exponent.

A confirmatory measure in the frequency domain that doesn’t depend on the assumption of scale-free activity was also used. MF was operationalized as the frequency that divides smoothed PSD in two equal halves [**Schwilden1985, Schwender1996, Golesorkhi2020**, 6, 31, 38, 47, 72, 74].

#### 2.4.2 Autocorrelation Window

The autocorrelation function (ACF) calculates the correlation between a time series and its lagged version using the following formula

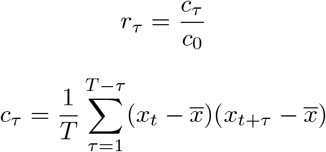

Where *r*_*τ*_ is autocorrelation at lag *τ*, 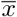 is time series, *x* is it’s mean, *T* is the number of time points and *c*_0_ is the sample variance of time series. Since our data did not contain any missing values, ACF was calculated by taking the inverse Fourier transform of the power spectral density, leveraging the Wiener-Khinchin theorem for faster calculations [1, 2]. Thus, ACW can be categorized with PLE and MF, connecting all three to the frequency domain. ACW-0 is defined as the first lag where autocorrelation function reaches zero [25, 60, 62, 67].

#### 2.4.3 Sample Entropy

SE was introduced by [7] as a variant of approximate entropy with two important advantages over it: independence with the length of the data and reduction of the bias caused by self-matching [7, 30, 65, 69, 72]. Given a time series *X* of length *N*, its SE for a pattern length *m* and similarity criterion *r* is computed as the negative natural logarithm of the probability that if two simultaneous data points of a subset *X*_*m*_ of length m have distance *d <*= *r* from each other, then two simultaneous data points of a subset *X*_*m*+1_ of length *m* + 1 also have distance *d <*= *r*.

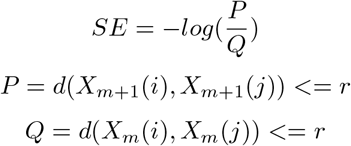

where *d*(*x, y*) is Chebyshev distance between *x* and *y*. For this paper, we set m as 2 and r as 0.5 following previous studies [30, 69].

#### 2.4.4 Lempel-Ziv Complexity

To calculate the LZC, each signal was converted into a binary sequence. Following [8, 11, 74], we used the median of the signal’s amplitude as the binarization threshold. After binarizing each region’s signal and converting it to a string sequence, the Lempel–Ziv algorithm[3] was used to compute LZC. As earlier studies have pointed out, LZC is dependent on the length of the sequence, thus a normalization factor (more info in [11]) was used to remove that effect. Briefly, the sequence is scanned from left to right and the LZC value was increased every time a new subsequence was encountered.

### 2.5 Gradient Index

To calculate the increase or decrease of measures in layers of self, we defined a measure called the gradient index. After seeing the hierarchical change of different measures across three layers of self, we calculated this rate of change by applying linear regression between the hierarchical positions (Internal = 1, External = 2, Mental = 3) and the measures at those positions. The slope of the best-fit line was defined as the gradient index for that variable. This procedure was applied to every measure for every scan, resulting in one gradient index per subject per measure per scan. A demonstration of the method is in figure 4 for Shanghai data. The same format for the UCLA dataset can be found in supplementary figure 4.

### 2.6 Network Analysis of Measures

To investigate the relationship between our different measures, we used measures from graph theory. Each measure was assigned to a node and connections between nodes were regularized partial correlations using Extended Bayesian Information Criterion and graphical least absolute shrinkage and selection operator (EBICglasso) method [14, 24, 46, 70]. We used two measures to quantify the position of each node in the network: strength of each node is the sum of the absolute input weights of that node whereas closeness is the inverse of the sum of all shortest paths from the node of interest to all other nodes. 10000 bootstraps were used to calculate confidence intervals of graph measures. The EBICglasso method relies on the assumption that the network is a sparse network. We also used the partial correlation method (non-regularized Gaussian Markov random field, pcor) that doesn’t depend on the assumption of sparsity. The results for pcor method can be found in supplementary figure 6.

The calculation of the measures have the possibility of biasing the network analysis. To explicate, both PLE and MF are calculated using the powerspectrum, and the ACW calculated from the autocorrelation function, which is the inverse Fourier transform of the power spectral density. High correlation and consequently, strength and closeness is expected from these measures regardless of physiological relevance. To exclude this possibility, we did the same analyses on simulated time series. We generated 1000 instances of white and pink noise of 250 time points and calculated the same measures via the same way on these simulated signals. Consequently, we did the network analyses on these simulated signals.

## 3 Results

### 3.1 Resting state: Dynamics of Self converge with its topographic hierarchy

Our first question was to investigate the differential dynamics of three self layers in the resting state. A non-parametric Kruskal-Wallis test was used to investigate the differences for each measure in both datasets. All tests were highly significant in both datasets (*p <* 0.001). Detailed results can be seen in supplementary results table S1 and S3 for Shanghai and UCLA respectively. Results of pairwise comparisons using Wilcoxon tests can be seen in figure 1, and the effect sizes are in supplementary tables S2 and S4.

**Figure 1:**
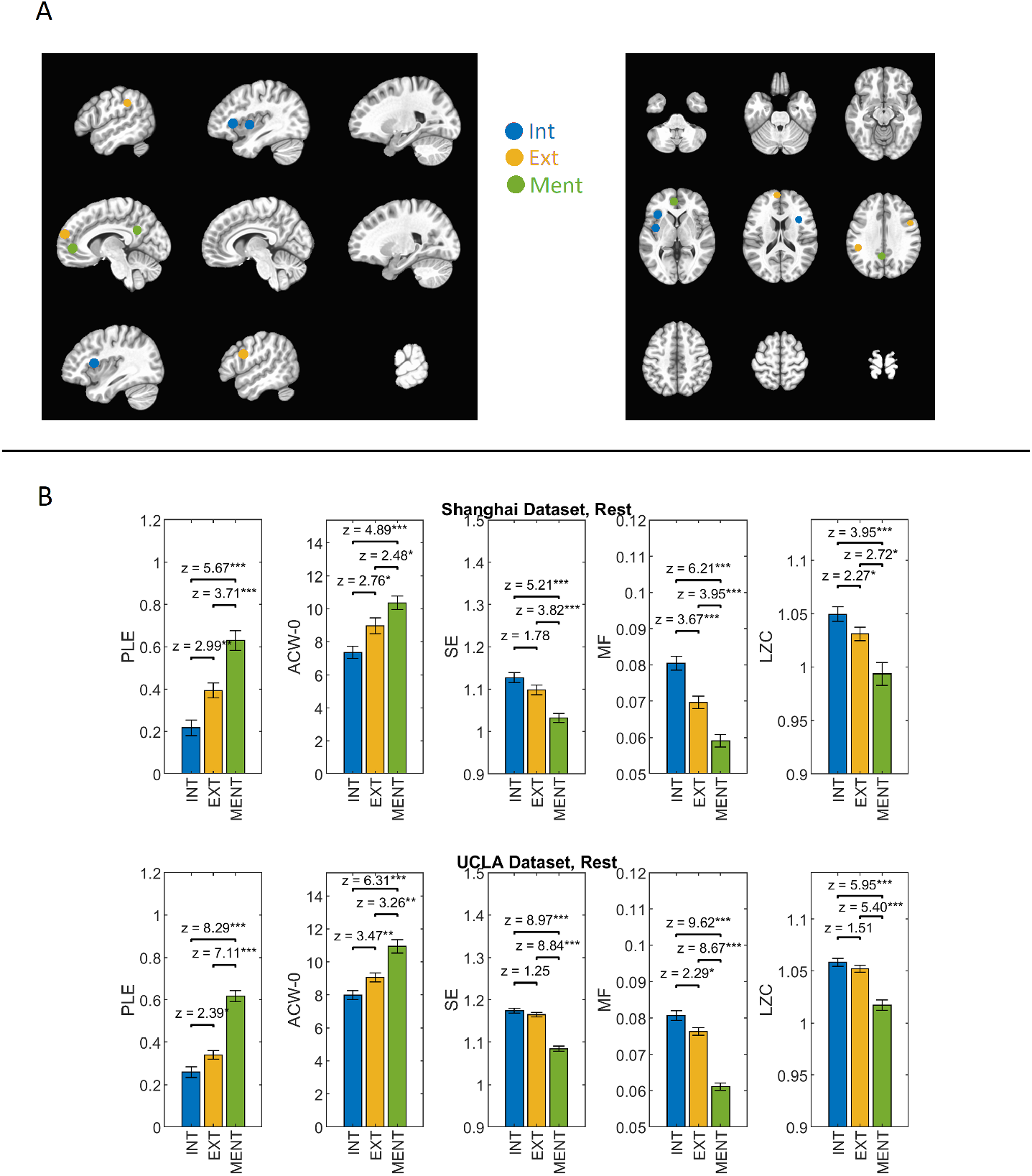
A. Interoceptive, Exteroceptive and Mental ROIs taken from [59]. BOLD signal of voxels inside 8 mm diameter spheres were averaged to get one time-series per one ROI (see table 1 for coordinates). After the calculation of measures, they were averaged to get one measure per layer per subject. B. Differences between layers calculated with pairwise Wilcoxon tests. Asterisks denote significance.

Our results show that all our measures (PLE, MF, LZC, ACW, SE) showed significant differences between the three layers of self. The mental layer shows the highest PLE, longest ACW, slowest MF, lowest LZC and lowest SE. Whereas the interoceptive layer exhibits the opposite pattern in all dynamic measures while the external layer is intermediate between the two other layers.

Together, these results suggest differentiation of the different layers of self according to their dynamics. Moreover, their topographical hierarchy converges with their dynamic hierarchy which follows the former.

### 3.2 Task-unspecific effects – carry-over of dynamic differentiation from rest to task

Next, we looked at task data to see the effect of tasks on dynamics in self layers. As can be seen in figure 2, this differentiation was carried over from rest to task. All the Kruskal-Wallis tests were highly significant (*p <* 0.001), details can be seen in supplementary results tables 5, 6, 7 and 8. Pairwise Wilcoxon tests of Pamenc, Stopsignal and Taskswitch be seen in figure 2. Results for other tasks can be found in supplementary figure 1. Of note, the *χ*^2^ values in the Shanghai task are lower than tasks in the UCLA dataset, with more cognitive tasks.

**Figure 2:**
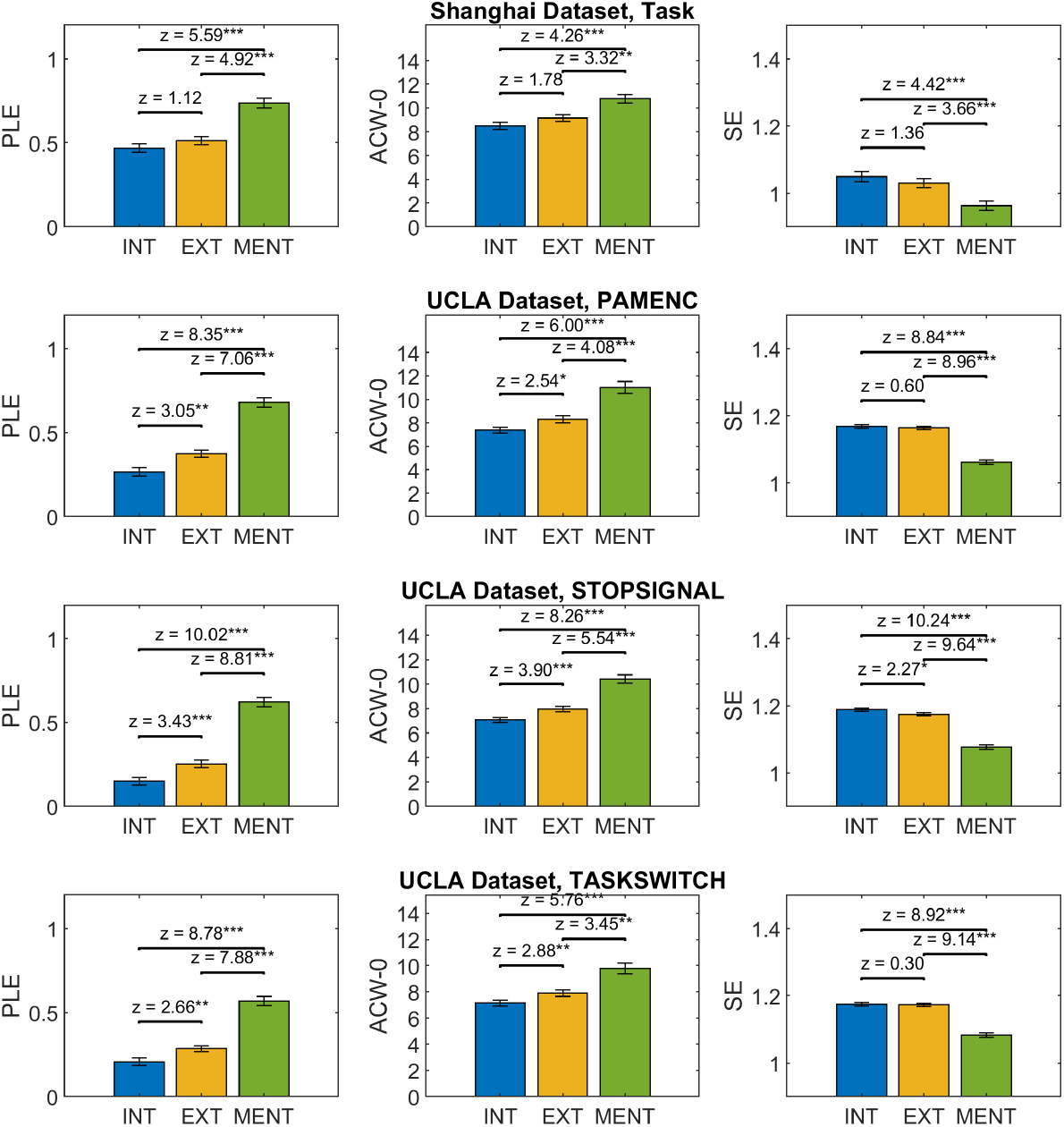
Pairwise Wilcoxon tests between three self layers. Asterisks denote significance.

Together, we observed similar topographic differentiation of the different measures in all task states of both data sets similar to their resting state. Hence, the convergence of topography and dynamics in the three layers of self is also manifested during the task. This suggests that the resting state topography is carried over to task states in a task-unspecific way – the topography and its dynamic are intrinsic to the brain as they are manifest in both rest and task states.

### 3.3 Rest-Task Difference of Dynamics is Sensitive to Task

Table 2. Rest - Task differences of the measures in three self layers. Plus and minus signs denote whether there is an increase or decrease from rest to task. Asterisks denote significance. Only significant differences were shown in the table.

Are there also task-specific effects during the transition from rest to task for the dynamics of layers? For that, we compared rest-task differences in all three layers across all our datasets using Wilcoxon tests. Results for PLE, ACW-0 and SE can be seen in figure 3, results for the remaining measures can be found in supplementary figures 2 and 3. A summary of significant differences can be found in table 2. Effect sizes for all the tests can be found in supplementary tables 9 and 10.

**Table 2:** Rest - Task Differences

| Measures | ROIs | Bart | Pamenc | Pamret | Scap | Stopsignal | Taskswitch | Shanghai |
| --- | --- | --- | --- | --- | --- | --- | --- | --- |
| PLE | INT |  |  |  |  | _** |  | +*** |
|  | EXT |  |  |  |  | _* |  | +* |
|  | MEN |  |  | _** | +** |  |  |  |
| ACW-0 | INT |  |  | _* |  | _* | _* | +** |
|  | EXT |  |  | _*** |  | _* | _** |  |
|  | MEN |  |  | _*** |  |  |  |  |
| MF | INT |  |  |  |  | + | +* | _*** |
|  | EXT |  |  | +*** |  |  | +** |  |
|  | MEN |  |  | +*** |  |  |  |  |
| SE | INT |  |  |  |  |  |  | _*** |
|  | EXT |  |  |  |  |  |  | _*** |
|  | MEN | _* | _* |  | _*** |  |  | _*** |
| LZC | INT | _*** | _*** | _*** | _*** |  | _* | _*** |
|  | EXT | _*** | _** | _*** | _*** |  |  | _*** |
|  | MEN | _*** | _** |  | _*** | _** |  | _** |

**Figure 3:**
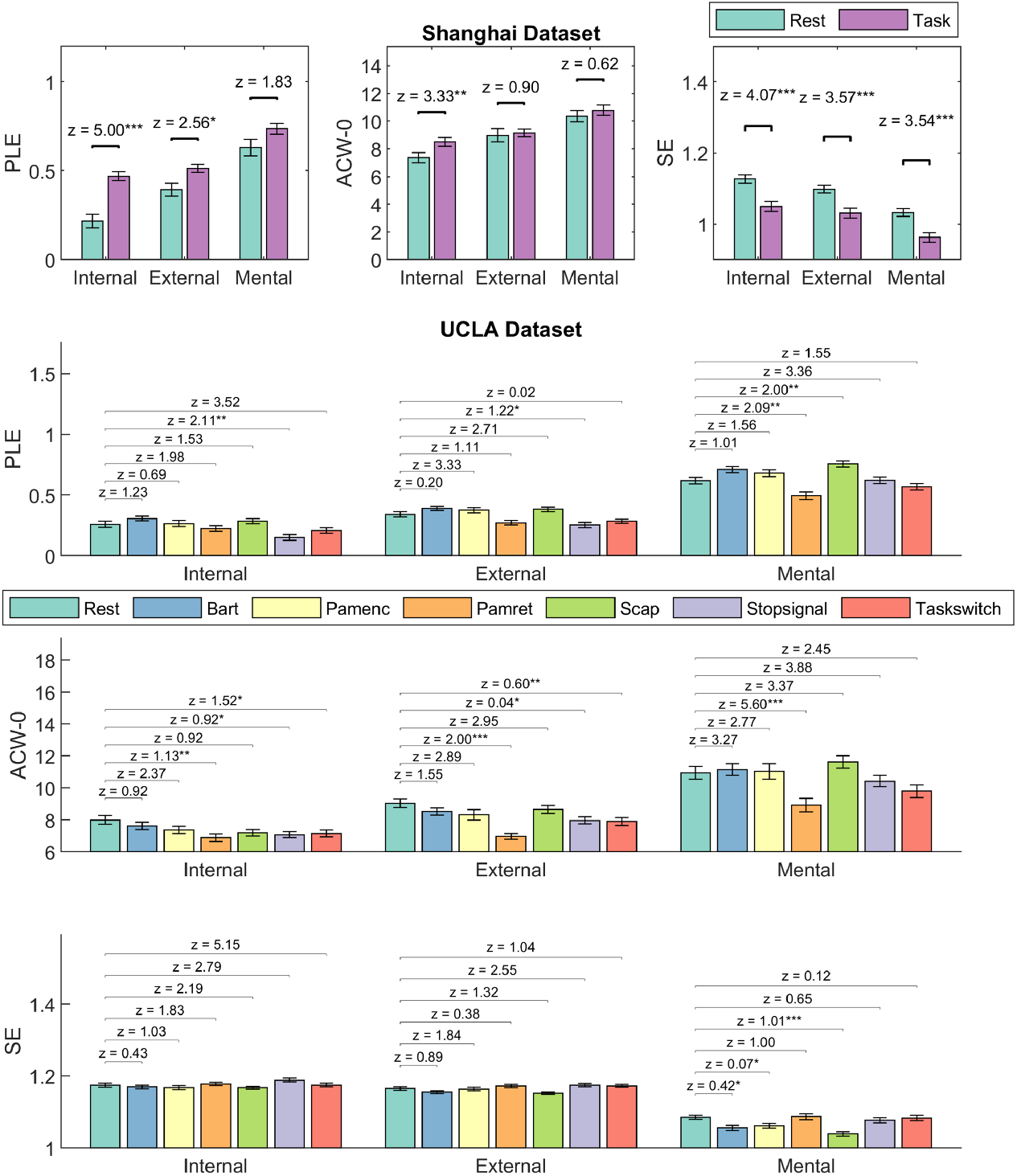
Comparison of measures in rest and task using Wilcoxon tests. Asterisks denote significance.

The internal self layer is the most sensitive one for the Shanghai task with the internally oriented listening-to-heartbeat task. Interestingly, the measures probing long-range temporal correlations (PLE, ACW-0, MF) were specific to the internal layer (increased PLE and ACW-0 and decreased MF from rest to task) whereas information related measures (SE and LZC) were layer-unspecific (decrease from rest to task in all layers).

On four of the seven UCLA tasks (Bart, Pamenc, Pamret, Scap), LZC showed a rather layer-unspecific effect (decrease in internal and external layers, also decrease in mental layer except for Pamret). Surprisingly, SE didn’t follow LZC, showing a decrease in only the mental layer in Bart, Pamenc and Scap.

PLE, MF and ACW-0, on the other hand, showed more task-specificity. A decrease in PLE and ACW-0 and corresponding increase in MF were observed in Pamret, Stopsignal and Taskswitch. PLE increase was observed in only the mental layer in Scap. PLE was decreased in the mental layer in Pamret; internal and external layers in Stopsignal; ACW-0 was decreased in all three layers in Pamret, and internal and external in Stopsignal and Taskswitch; MF was increased in external and mental layers in Pamret and internal and external layers in Taskswitch. Two things of note. Firstly, while they’re both memory-related tasks, Pamret caused significant changes in our measures whereas Pamenc did not cause any. The reasons for this will be discussed in the discussion. Secondly, while information-related measures showed some change in at least one layer in all tasks, temporal measures didn’t show any change in two of UCLA tasks.

### 3.4 Topographic hierarchy of self: Task-specific and unspecific changes

To measure the degree of hierarchy among the three self-layers, we devised the new measure gradient index (see Methods). The main question here was whether the degree of hierarchy changed during the transition from rest to task states and whether that occurred in a task-specific or -unspecific way. For that, we measured the gradient index in all recordings for all subjects. The differences in gradient indices between rest and task were tested with Wilcoxon tests. Results for comparisons can be seen in figure 4. Effect sizes can be found in supplementary tables 11 and 12.

**Figure 4:**
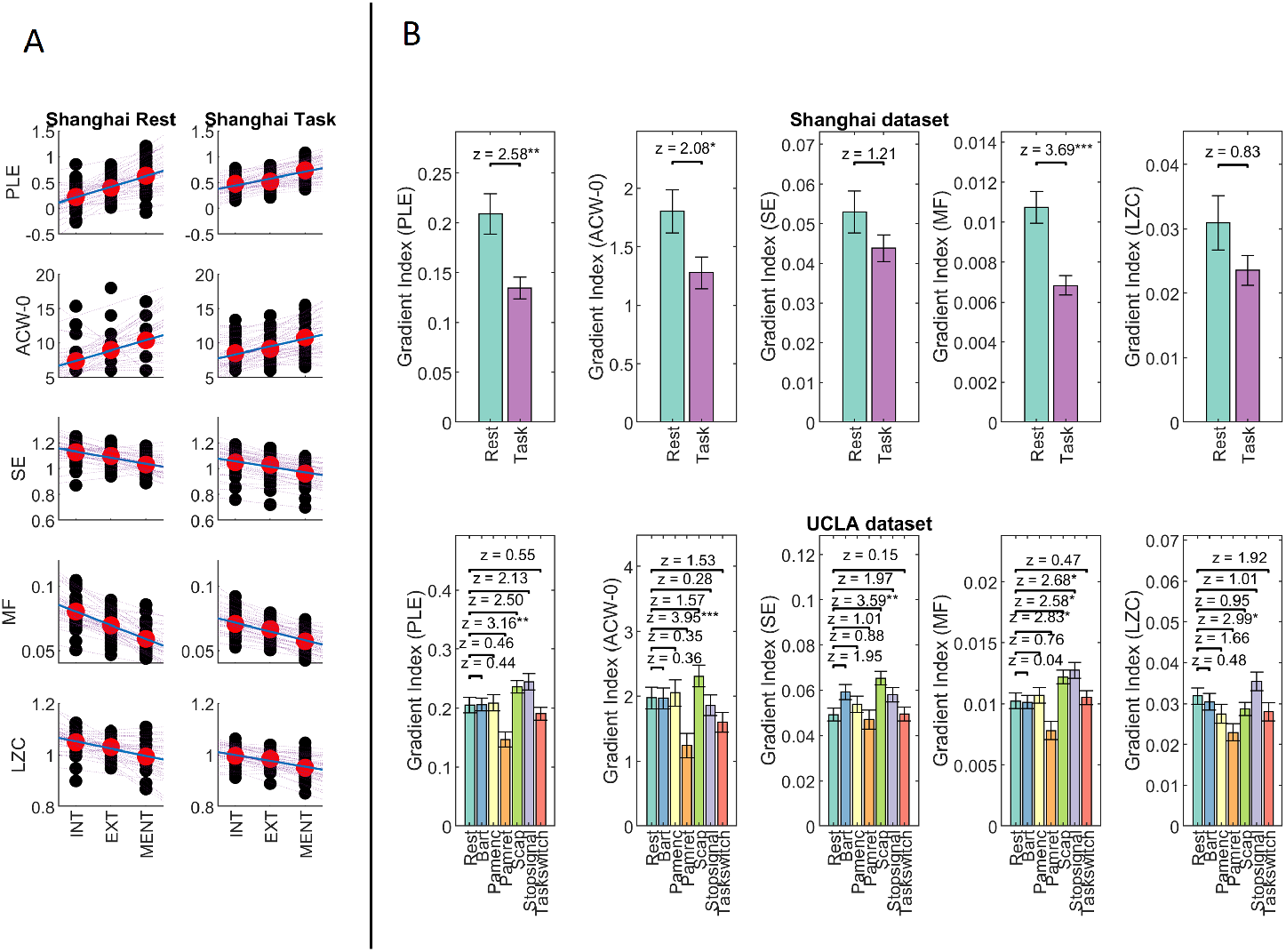
A. Calculation of the gradient index. A best-fit linear regression line between the measures calculated in three layers was calculated for each measure in each subject in order to probe the hierarchic structure of self layers. The gradient index was defined as the slope of the best-fit line. B. Comparison of gradient indices in rest and task using Wilcoxon tests.

The results indicate that the degree of hierarchy changes in those measures that reflect long-range temporal connections (PLE, MF and ACW-0) and respond to the internal task of the Shanghai dataset. In contrast, that was not the case in the measures based on information processing (SE and LZC) in the case of the Shanghai dataset. In the UCLA dataset, once again Pamret seems to be close to the Shanghai dataset showing decreased gradient indices of PLE, MF and ACW; and also LZC. The gradient index based on MF also responded to Scap and Stopsignal. Surprisingly gradient index of SE increased from rest to task in Scap.

Together, albeit tentatively, the results indicate differential roles of information-related and temporal measures. More importantly, we showed that different tasks affect the dynamic topography of self in slightly distinct ways with the most prominent example being the internal Shanghai task and the self-specific Pamenc UCLA task. More generally, our results show the malleability of the hierarchical topography of self with both task-specific and -unspecific effects.

### 3.5 Hierarchy of dynamics and its malleability during task states

We observed that the topographic three-layer hierarchy of self changes during task states in both task-specific and-unspecific ways. We also observed changes in different dynamic measures during task states. Does this mean that there is a hierarchy among the dynamic measures? In order to probe such a dynamic hierarchy, we calculated two graph-theory measures, strength and closeness in the networks based on our 5 measures. Results for the Shanghai dataset and Pamenc, Stopsignal and Taskswitch of UCLA can be seen in figure 5. Results for other UCLA tasks are in supplementary figure 5. Additionally, we replicated the analysis using the pcor method that doesn’t depend on the assumption of sparsity that’s inherent to EBICglasso and found the same results. The results for the pcor method can be found in supplementary figure 6.

**Figure 5:**
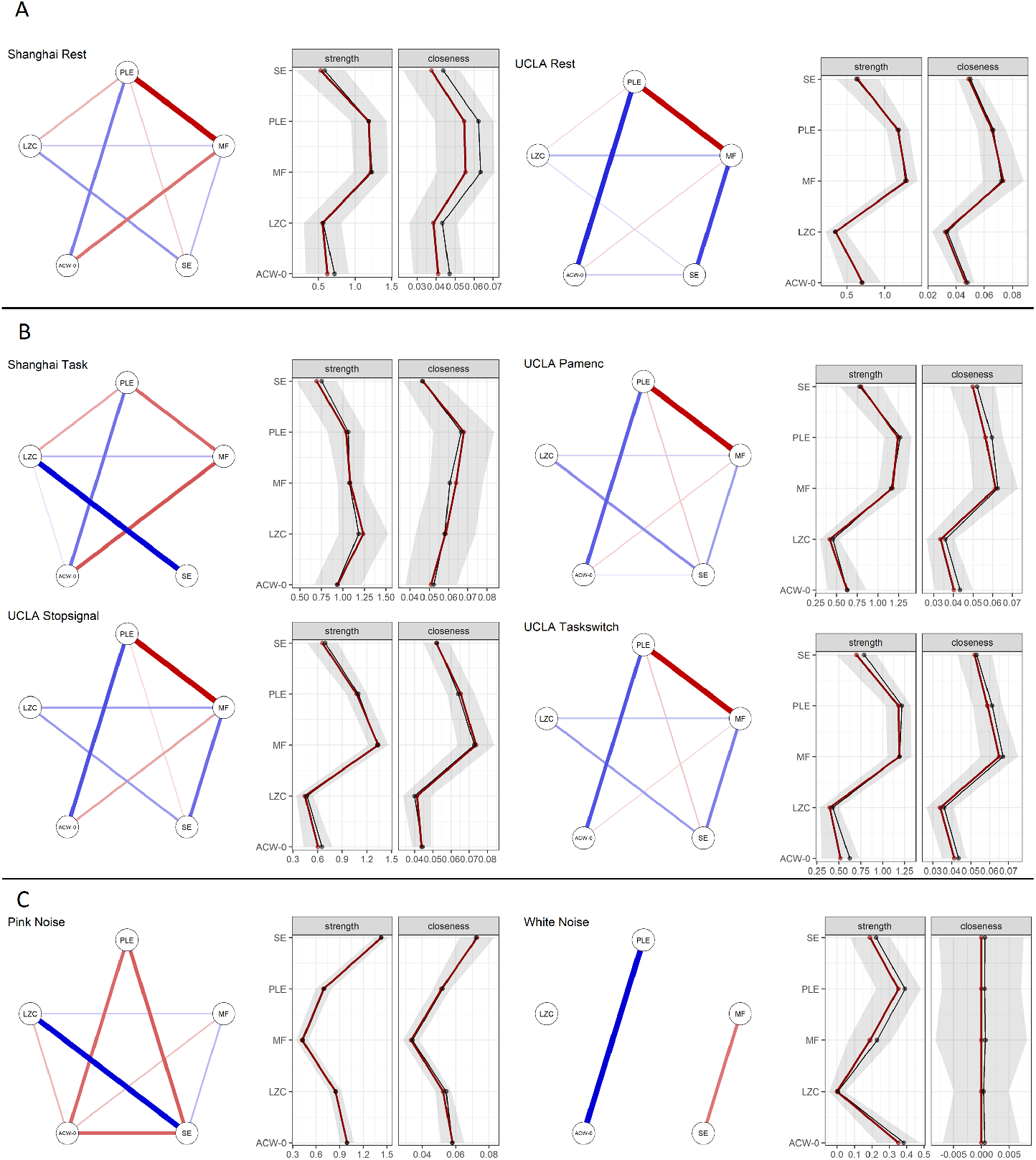
Network analysis of dynamic measures. The network structure between different measures was calculated with the EBICglasso method. Strength and closeness were calculated from the networks. In network plots, red lines show negative relations whereas blue lines show positive ones. Line thickness shows the degree of relation. In the line plots, gray areas indicate confidence intervals using bootstrapping 10000 samples. Black lines show bootstrap estimates of network measures whereas red ones show network estimates from the original sample. A. Network analysis in rest. B. Same analyses in the task states. C. Network analyses on simulated signals.

In the resting state, PLE and MF showed the highest strength and closeness in both datasets, followed by ACW-0, SE and LZC. However, this relationship changed in favor of SE and LZC in the Shanghai task, whereas all UCLA datasets kept the same relationship between the measures during the different tasks. This suggests that the relationship between the different measures is by itself prone to change e.g. being dynamic by itself showing a certain degree of task-specificity.

To exclude the possibility of bias stemming from the calculation of these measures, we did the same analyses on simulated signals (see methods). Network analyses on 1000 instances of white noise and 1000 instances of pink noise which has similar scale-free properties are presented in figure 5C. On pink noise, we see a different structure of measures with low PLE and MF and high SE and ACW-0 strength. On white noise, the structure between strength of measures is preserved, however, the strength is considerably lower. Closeness is interestingly, 0. The same observations were made using the pcor method which is presented on supplementary figure 7. These simulation results suggest that the network results we found have a physiological basis in the brain signal rather than being due to the intrinsic (e.g. mathematical) properties of these measures themselves.

## 4 Discussion

The aim of this paper is twofold: First, we investigate the dynamic features of the intrinsic topography of a higher-order cognitive function like the self. Second, we are interested in how these dynamics change with different tasks including task-specific and -unspecific effects. We used two different fMRI datasets with a total of two resting states and seven different tasks to probe these two questions. We extracted five different measures: PLE, MF and ACW-0, probing dynamics through long-range temporal correlations and structure of powerspectrum; SE and LZC for information complexity and predictability of the BOLD signal.

### 4.1 Dynamics follows the topographic hierarchy of the three layers of self

A previous meta-analysis by [59] shows a nested structure of self regions with the mental layer encompassing external and internal layers; and external layer encompassing internal layer. We therefore hypothesize a hierarchy of dynamics in self regions, following the hierarchy of intrinsic neural timescales in global [42, 48, 67, 71, 74] and local (see [62, 72] for a case of sensory input regions) levels (see supplementary figure 1 for an overlay of the ROIs in this paper and transmodal networks as determined in [55]). We found increasing PLE and ACW-0 from internal to mental layers and a corresponding decrease in MF in the resting state. This was complemented by decreasing SE and LZC. These results are in line with the hypotheses surrounding longer and shorter neural timescales suitable for temporal integration and segregation, respectively. High power in slow frequencies [67, 74, 83] provide longer temporal receptive windows as a basis for summing and pooling of different stimuli and vica versa [16, 33–35, 42, 48, 67, 83]. The decreasing of LZC and SE is in accordance with a previous study [74], though implications are tentative, this might be the result of the aforementioned pooling. Temporally smoothed activity is more orderly and predictable than the activity with higher complexity, which is suitable for segregation [83].

Our findings extend the previous data. The hierarchical organization of self is based on a meta-analysis of separate task studies on self [59]. We here demonstrate that an analogous hierarchy can be observed in both the resting state and task states. This strongly suggests the intrinsic nature of the hierarchical topography of self as it is not affected as such in its basic organization but only modulated (in small degrees, if at all) by the task states. In other terms, due to its intrinsic nature, the topographic hierarchy of self with its three layers is carried over from rest to task states.

Importantly, we observed the hierarchy of self in both our dynamic and information processing measures. The topographic hierarchy of self with its three layers thus converges with a corresponding dynamic hierarchy. This strongly suggests the intrinsic nature of the convergence of topography and hierarchy for a higher-order cognitive function like the self that extends beyond previous findings that show an intrinsic topographic organization of the brain itself like core-periphery distinction [55, 60, 67]. Our findings extend the existence of such intrinsic topographic organization from the brain to higher-order cognitive functions like self: they are intrinsically ingrained within the brain’s topography and dynamics even already in the brain’s spontaneous activity itself prior to and independent of any external stimuli or specific cognition during task-related activity. Hence, higher-order cognitive functions like self follow the basic organizational principles of the brain itself, in their topographic hierarchical organization and its convergence with dynamics.

### 4.2 Task-specific and -unspecific effects in both topography and dynamics

In a further step, we compared dynamics in rest and task states. We found that the change of dynamics shows task and layer specificity, the different layers do not respond to each task in the same way nor do they respond the same in every layer. As can be seen in table 2, the dynamic measures PLE, MF and ACW-0 showed more layer and task-specific effects compared to information measures LZC and SE. This indicates that different regions are defined by their temporal structure in the way they respond to external perturbations. LZC and SE on the other hand are more task and layer unspecific, thus, more malleable measures for external perturbation, further supporting their role as foreground measures [82].

The difference between Pamenc and Pamret is an intriguing case. While both are memory-related tasks, there are some subtle psychological differences. Pamret focuses on memory retrieval and episodic simulation thus involving a high degree of self-specificity while this is distinct in the encoding task probed in Pamenc. Only Pamret showed significant differences from rest to task with seemingly task-specific effects in the dynamic measures of the three layers. On the other hand, Pamenc, a memorization task focusing on external stimuli, showed no difference in any of the temporal measures. The reason for this, we claim, is the nature of the topography. We here probe the topography of self; this let us assume that neural activity in these regions may be particularly sensitive to self-specific tasks like the Pamret and our interoceptive awareness task (with both tasks showing more or less similar task-related changes). Albeit tentatively, we therefore suppose that task-specific effects in both topographic layers and dynamic measures are more likely to be associated with self-specific stimuli or tasks. While tasks or stimuli that involve to a lower degree the self, i..e, non-self-specific, may rather show task-unspecific effects in the dynamics of the topography of self.

To probe the relationship between task-related changes and topography further, we devised a new measure called gradient index (see Methods). Gradient index measures the hierarchical structure of dynamics i.e. how much change for any measure do we see when we go from internal to external to mental. To see the effect of different tasks on this gradient, we compared gradient indices at task and rest. In the Shanghai task, we saw a significant decrease in gradient index from rest to task. We saw the same effect on Pamret too, with also an increase in LZC. The Pamenc - Pamret distinction can here too be as a further confirmation of the findings we discussed in the above paragraph.

### 4.3 Relationship Between Different Measures is Hierarchical and Changes With Task

Changing our focus from topography to dynamics themselves, we investigated the relationship between different measures using a graph-theoretic network approach. In both datasets, we, during the resting state, saw a hierarchy of measures with temporal-dynamic measures PLE and MF representing a hub with high strength and closeness and ACW-0 being close while LZC and SE showed the smallest values. Importantly, we controlled for the physiological (rather than mathematical or otherwise) nature of these dynamic relationships by conducting the network analysis on simulated data and finding incompatible results.

Together, these data suggest hierarchical organization among the dynamic measures themselves. Dynamic measures like ACW and PLE operate as dynamic core while information processing (SE, LZC) takes on the role of dynamic periphery. Although preliminary, we therefore suppose that there may be analogous forms of hierarchical organizations, e.g., core-periphery, in both spatial-topographic and temporal-dynamic domains.

This is further supported by the task data. This relationship among the different measures was more or less preserved during and thus carried over to the task states in the UCLA data. In contrast, the dynamic relationship among different measures changed in the self-specific task of the Shanghai data set. This reveals the malleability of the relationship between the dynamic and information processing measures. Specifically, the PLE, ACW and MF remained relatively stable in their closeness and strength while SE and LZC showed stronger changes. That does not only compare well with the core-periphery organization on a dynamic level but also with the recently supposed distinction of stable background dynamic measures and more flexible foreground dynamic measures [82].

## 5 Conclusion

Using two datasets including rest and seven task states, we investigated whether a higher-order cognitive function like self is characterized by an intrinsic topographic-dynamic hierarchy within the brain’s inner core during both rest and task states.

We observed that various dynamic and information processing measures followed the topographic hierarchy of self during both rest and task states. Subtracting task from rest states yielded task-unspecific and -specific tasks in both topographic and dynamic measures. Finally, we also demonstrate a hierarchy among the various measures themselves. Dynamic measures (like PLE) are stable during both rest and task thus remaining in the background while information measures (SE, LZC) are more flexible, operating in the foreground.

Together, we demonstrate an intrinsic topography and its convergence with dynamics for a higher-order cognitive function like the self within the brain’s core. This does not only support the intrinsic nature of topography and dynamic on the neural level of the brain but also its intrinsic nature for higher-order cognitive functions like self. Higher-order cognitive functions may thus by themselves be characterized in spatial-topographic and temporal-dynamic terms, just as postulated by the recently proposed concept of Spatiotemporal Neuroscience [57, 58]. Topography and dynamic may then be shared by both neural and cognitive/psychological levels as their “common currency”.

## Acknowledgements

This work was supported by EJLB-Michael Smith Foundation; Canadian Institutes of Health Research; Ministry of Science and Technology of China; National Science Foundation of China (Grant/Award Numbers: 82101582); National Key R&D Program of China (2016YFC1306700); Hope of Depression Foundation (HDRF); Start-Up Research Grant in Hangzhou Normal University and European Union’s Horizon 2020 Framework Program for Research; Innovation under the Specific Grant Agreement No. 785907 (Human Brain Project SGA2) and Canada-UK Artificial Intelligence (AI) Initiative “The self as agentenvironment nexus: crossing disciplinary boundaries to help human selves and anticipate artificial selves” (ES/T01279X/1) (together with Karl J. Friston from the UK).

## Author Contributions

Conceptualization: Y.Ç., G.N.; Methodology: Y.Ç., G.N.; Software: Y.Ç.; Formal Analysis: Y.Ç., M.A.G.; Visualization: Y.Ç.; Funding acquisition: G.N.; Data acquisition: C.L., J.W., H.C., W.L.; Supervision: G.N.; Writing: Y.Ç., G.N.

## References

1. Wiener, N. Generalized harmonic analysis. Acta Mathematica 55, 117–258. https://doi.org/10.1007/BF02546511 (Jan. 1930).

2. Khintchine, A. Korrelationstheorie der stationären stochastischen Prozesse. Mathematische Annalen 109, 604–615. ISSN: 1432-1807. https://doi.org/10.1007/BF01449156 (1934).

3. Lempel, A. & Ziv, J. On the Complexity of Finite Sequences. IEEE Transactions on Information Theory 22, 75–81. ISSN: 1557-9654 VO-22 (1976).

4. Auger, F. & Flandrin, P. Improving the readability of time-frequency and time-scale representations by the reassignment method. IEEE Transactions on Signal Processing 43, 1068–1089 (1995).

5. Cox, R. W. AFNI: Software for Analysis and Visualization of Functional Magnetic Resonance Neuroimages. Computers and Biomedical Research 29, 162–173. ISSN: 0010-4809. https://www.sciencedirect.com/science/article/pii/S0010480996900142 (1996).

6. McDonald, T., Berkowitz, R. & Hoffman, W. E. Median EEG Frequency is More Sensitive to Increases in Sympathetic Activity Than Bispectral Index. Journal of Neurosurgical Anesthesiology 11. ISSN: 0898-4921. https://journals.lww.com/jnsa/Fulltext/1999/10000/Median_EEG_Frequency_is_More_Sensitive_to.5.aspx (1999).

7. Richman, J. & Moorman, J. Physiological Time-Series Analysis Using Approximate Entropy and Sample Entropy. American journal of physiology. Heart and circulatory physiology 278, H2039–49 (July 2000).

8. Nagarajan, R. Quantifying physiological data with Lempel-Ziv complexity - Certain issues. IEEE transactions on bio-medical engineering 49, 1371–1373 (Dec. 2002).

9. Critchley, H. D., Wiens, S., Rotshtein, P., Ohman, A. & Dolan, R. J. Neural systems supporting interoceptive awareness. eng. Nature neuroscience 7, 189–195. ISSN: 1097-6256 (Print) (Feb. 2004).

10. Northoff, G. & Bermpohl, F. Cortical midline structures and the self. eng. Trends in cognitive sciences 8, 102–107. ISSN: 1364-6613 (Print) (Mar. 2004).

11. Aboy, M., Hornero, R., Abásolo, D. & Alvarez, D. Interpretation of the Lempel-Ziv Complexity Measure in the Context of Biomedical Signal Analysis. IEEE transactions on bio-medical engineering 53, 2282–2288 (Dec. 2006).

12. Fulop, S. A. & Fitz, K. Algorithms for computing the time-corrected instantaneous frequency (reassigned) spectrogram, with applications. The Journal of the Acoustical Society of America 119, 360–371. ISSN: 0001-4966. https://doi.org/10.1121/1.2133000 (Jan. 2006).

13. Northoff, G. et al. Self-referential processing in our brain–a meta-analysis of imaging studies on the self. eng. NeuroImage 31, 440–457. ISSN: 1053-8119 (Print) (May 2006).

14. Friedman, J., Hastie, T. & Tibshirani, R. Sparse inverse covariance estimation with the graphical lasso. Biostatistics 9, 432–441. ISSN: 1465-4644. https://doi.org/10.1093/biostatistics/kxm045 (2007).

15. Pollatos, O., Schandry, R., Auer, D. P. & Kaufmann, C. Brain structures mediating cardiovascular arousal and interoceptive awareness. eng. Brain research 1141, 178–187. ISSN: 0006-8993 (Print) (Apr. 2007).

16. Kiebel, S. J., Daunizeau, J. & Friston, K. J. A Hierarchy of Time-Scales and the Brain. PLOS Computational Biology 4, e1000209. https://doi.org/10.1371/journal.pcbi.1000209 (Nov. 2008).

17. Northoff, G. & Panksepp, J. The trans-species concept of self and the subcortical-cortical midline system. eng. Trends in cognitive sciences 12, 259–264. ISSN: 1364-6613 (Print) (July 2008).

18. He, B. J., Zempel, J. M., Snyder, A. Z. & Raichle, M. E. The temporal structures and functional significance of scale-free brain activity. eng. Neuron 66, 353–369. ISSN: 1097-4199. https://pubmed.ncbi.nlm.nih.gov/20471349#20https://www.ncbi.nlm.nih.gov/pmc/articles/PMC2878725/ (May 2010).

19. Jo, H. J., Saad, Z. S., Simmons, W. K., Milbury, L. A. & Cox, R. W. Mapping sources of correlation in resting state FMRI, with artifact detection and removal. NeuroImage 52, 571–582. ISSN: 1053-8119. https://www.sciencedirect.com/science/article/pii/S1053811910006580 (2010).

20. Qin, P. et al. Anterior cingulate activity and the self in disorders of consciousness. eng. Human brain mapping 31, 1993–2002. ISSN: 1097-0193 (Electronic) (Dec. 2010).

21. Van der Meer, L., Costafreda, S., Aleman, A. & David, A. S. Self-reflection and the brain: a theoretical review and meta-analysis of neuroimaging studies with implications for schizophrenia. eng. Neuroscience and biobehavioral reviews 34, 935–946. ISSN: 1873-7528 (Electronic) (May 2010).

22. Wiebking, C. et al. Abnormal body perception and neural activity in the insula in depression: An fMRI study of the depressed “material me”. The world journal of biological psychiatry : the official journal of the World Federation of Societies of Biological Psychiatry 11, 538–549 (2010).

23. Blanke, O. Multisensory brain mechanisms of bodily self-consciousness. Nature Reviews Neuroscience 13, 556–571. ISSN: 1471-0048. https://doi.org/10.1038/nrn3292 (2012).

24. Epskamp, S., Cramer, A., Waldorp, L., Schmittmann, V. & Borsboom, D. qgraph: Network Visualizations of Relationships in Psychometric Data. Journal of statistical software 48 (Apr. 2012).

25. Honey, C. J. et al. Slow cortical dynamics and the accumulation of information over long timescales. eng. Neuron 76, 423–434. ISSN: 1097-4199. https://pubmed.ncbi.nlm.nih.gov/23083743#20https://www.ncbi.nlm.nih.gov/pmc/articles/PMC3517908/ (Oct. 2012).

26. Schacter, D. L. et al. The future of memory: remembering, imagining, and the brain. eng. Neuron 76, 677–694. ISSN: 1097-4199 (Electronic) (Nov. 2012).

27. Gotts, S. J. et al. The perils of global signal regression for group comparisons: a case study of Autism Spectrum Disorders. eng. Frontiers in human neuroscience 7, 356. ISSN: 1662-5161. https://pubmed.ncbi.nlm.nih.gov/23874279#20https://www.ncbi.nlm.nih.gov/pmc/articles/PMC3709423/ (July 2013).

28. Prebble, S. C., Addis, D. R. & Tippett, L. J. Autobiographical memory and sense of self. eng. Psychological bulletin 139, 815–840. ISSN: 1939-1455 (Electronic) (July 2013).

29. Tagliazucchi, E. et al. Breakdown of long-range temporal dependence in default mode and attention networks during deep sleep. Proceedings of the National Academy of Sciences 110, 15419LP–15424. http://www.pnas.org/content/110/38/15419.abstract (Sept. 2013).

30. McDonough, I. M. & Nashiro, K. Network complexity as a measure of information processing across resting-state networks: evidence from the Human Connectome Project 2014. https://www.frontiersin.org/article/10.3389/fnhum.2014.00409.

31. Bachiller, A. et al. Decreased entropy modulation of EEG response to novelty and relevance in schizophrenia during a P300 task. European Archives of Psychiatry and Clinical Neuroscience 265, 525–535. ISSN: 1433-8491. https://doi.org/10.1007/s00406-014-0525-5 (2015).

32. Blanke, O., Slater, M. & Serino, A. Behavioral, Neural, and Computational Principles of Bodily Self-Consciousness. eng. Neuron 88, 145–166. ISSN: 1097-4199 (Electronic) (Oct. 2015).

33. Chaudhuri, R., Knoblauch, K., Gariel, M. A., Kennedy, H. & Wang, X. J. A Large-Scale Circuit Mechanism for Hierarchical Dynamical Processing in the Primate Cortex. Neuron 88, 419–431. ISSN: 10974199. http://dx.doi.org/10.1016/j.neuron.2015.09.008 (2015).

34. Gollo, L. L., Zalesky, A., Hutchison, R. M., van den Heuvel, M. & Breakspear, M. Dwelling quietly in the rich club: brain network determinants of slow cortical fluctuations. Philosophical Transactions of the Royal Society B: Biological Sciences 370, 20140165. https://doi.org/10.1098/rstb.2014.0165 (May 2015).

35. Murray, R. J., Debbané, M., Fox, P. T., Bzdok, D. & Eickhoff, S. B. Functional connectivity mapping of regions associated with self- and other-processing. eng. Human brain mapping 36, 1304–1324. ISSN: 1097-0193 (Electronic) (Apr. 2015).

36. Sui, J. & Humphreys, G. W. The Integrative Self: How Self-Reference Integrates Perception and Memory. eng. Trends in cognitive sciences 19, 719–728. ISSN: 1879-307X (Electronic) (Dec. 2015).

37. Taylor, P., Hobbs, J. N., Burroni, J. & Siegelmann, H. T. The global landscape of cognition: hierarchical aggregation as an organizational principle of human cortical networks and functions. Scientific Reports 5, 18112. ISSN: 2045-2322. https://doi.org/10.1038/srep18112 (2015).

38. Verrusio, W. et al. The Mozart Effect: A quantitative EEG study. Consciousness and Cognition 35, 150–155. ISSN: 1053-8100. https://www.sciencedirect.com/science/article/pii/S1053810015001130 (2015).

39. Wiebking, C. & Northoff, G. Neural activity during interoceptive awareness and its associations with alexithymia-An fMRI study in major depressive disorder and non-psychiatric controls. eng. Frontiers in psychology 6, 589. ISSN: 1664-1078 (Print) (2015).

40. Christoff, K., Irving, Z. C., Fox, K. C. R., Spreng, R. N. & Andrews-Hanna, J. R. Mind-wandering as spontaneous thought: a dynamic framework. Nature Reviews Neuroscience 17, 718–731. ISSN: 1471-0048. https://doi.org/10.1038/nrn.2016.113 (2016).

41. Huang, Z., Obara, N., Davis, H. H. 4., Pokorny, J. & Northoff, G. The temporal structure of resting-state brain activity in the medial prefrontal cortex predicts self-consciousness. eng. Neuropsychologia 82, 161–170. ISSN: 1873-3514 (Electronic) (Feb. 2016).

42. Margulies, D. S. et al. Situating the default-mode network along a principal gradient of macroscale cortical organization. Proceedings of the National Academy of Sciences 113, 12574LP–12579. http://www.pnas.org/content/113/44/12574.abstract (Nov. 2016).

43. Northoff, G. Is the self a higher-order or fundamental function of the brain? The “basis model of self-specificity” and its encoding by the brain’s spontaneous activity. eng. Cognitive neuroscience 7, 203–222. ISSN: 1758-8936 (Electronic) (2016).

44. Poldrack, R. A. et al. A phenome-wide examination of neural and cognitive function. Scientific Data 3, 160110. ISSN: 2052-4463. https://doi.org/10.1038/sdata.2016.110 (2016).

45. Gorgolewski, K. J., Durnez, J. & Poldrack, R. A. Preprocessed Consortium for Neuropsychiatric Phenomics dataset [version 2; peer review: 2 approved]. F1000Research 6. https://f1000research.com/articles/6-1262/v2 (2017).

46. Epskamp, S., Borsboom, D. & Fried, E. I. Estimating psychological networks and their accuracy: A tutorial paper. Behavior Research Methods 50, 195–212. ISSN: 1554-3528. https://doi.org/10.3758/s13428-017-0862-1 (2018).

47. Huang, Z., Liu, X., Mashour, G. A. & Hudetz, A. G. Timescales of Intrinsic BOLD Signal Dynamics and Functional Connectivity in Pharmacologic and Neuropathologic States of Unconsciousness. The Journal of Neuroscience 38, 2304LP–2317. http://www.jneurosci.org/content/38/9/2304.abstract (Feb. 2018).

48. Huntenburg, J. M., Bazin, P.-L. & Margulies, D. S. Large-Scale Gradients in Human Cortical Organization. Trends in Cognitive Sciences 22, 21–31. ISSN: 1364-6613. https://doi.org/10.1016/j.tics.2017.11.002 (Jan. 2018).

49. Lin, Y.-T. Visual Perspectives in Episodic Memory and the Sense of Self. eng. Frontiers in psychology 9, 2196. ISSN: 1664-1078 (Print) (2018).

50. Ji, J. L. et al. Mapping the human brain’s cortical-subcortical functional network organization. NeuroImage 185, 35–57. ISSN: 1053-8119. https://www.sciencedirect.com/science/article/pii/S1053811918319657 (2019).

51. Wolff, A. et al. The temporal signature of self: Temporal measures of resting-state EEG predict self-consciousness. eng. Human brain mapping 40, 789–803. ISSN: 1097-0193 (Electronic) (Feb. 2019).

52. Zhang, J., Huang, Z., Tumati, S. & Northoff, G. Intrinsic Architecture of Global Signal Topography and Its Modulation by Tasks. bioRxiv, 798819. https://www.biorxiv.org/content/10.1101/798819v1 (Jan. 2019).

53. Cui, H. et al. Insula shows abnormal task-evoked and resting-state activity in first-episode drug-näıve generalized anxiety disorder. eng. Depression and Anxiety 37, 632–644. ISSN: 15206394 (July 2020).

54. Frewen, P. et al. Neuroimaging the consciousness of self: Review, and conceptual-methodological framework. Frewen, Paul, 2020.

55. Ito, T., Hearne, L. J. & Cole, M. W. A cortical hierarchy of localized and distributed processes revealed via dissociation of task activations, connectivity changes, and intrinsic timescales. NeuroImage 221, 117141. ISSN: 1053-8119 (Nov. 2020).

56. Kolvoort, I. R., Wainio-Theberge, S., Wolff, A. & Northoff, G. Temporal integration as “common currency” of brain and self-scale-free activity in resting-state EEG correlates with temporal delay effects on self relatedness. Human Brain Mapping 41, 4355–4374. https://onlinelibrary.wiley.com/doi/abs/10.1002/hbm.25129 (2020).

57. Northoff, G., Wainio-Theberge, S. & Evers, K. Is temporo-spatial dynamics the “common currency” of brain and mind? In Quest of “Spatiotemporal Neuroscience”. Physics of Life Reviews 33, 34–54. ISSN: 1571-0645. https://www.sciencedirect.com/science/article/pii/S1571064519300739 (2020).

58. Northoff, G., Wainio-Theberge, S. & Evers, K. Spatiotemporal neuroscience – what is it and why we need it. Physics of Life Reviews 33, 78–87. ISSN: 1571-0645. https://www.sciencedirect.com/science/article/pii/S157106452030035X (2020).

59. Qin, P., Wang, M. & Northoff, G. Linking bodily, environmental and mental states in the self-A three-level model based on a meta-analysis. eng. Neuroscience and biobehavioral reviews 115, 77–95. ISSN: 1873-7528 (Electronic) (Aug. 2020).

60. Raut, R. V., Snyder, A. Z. & Raichle, M. E. Hierarchical dynamics as a macroscopic organizing principle of the human brain. Proceedings of the National Academy of Sciences 117, 20890LP–20897. http://www.pnas.org/content/117/34/20890.abstract (Aug. 2020).

61. Scalabrini, A. et al. All roads lead to the default-mode network—global source of DMN abnormalities in major depressive disorder. Neuropsychopharmacology 45, 2058–2069. ISSN: 1740-634X. https://doi.org/10.1038/s41386-020-0785-x (2020).

62. Wengler, K., Goldberg, A. T., Chahine, G. & Horga, G. Distinct hierarchical alterations of intrinsic neural timescales account for different manifestations of psychosis. eLife 9 (eds Frank, M. J., Gillan, C. M. & Corlett, P. R.) e56151. ISSN: 2050-084X. https://doi.org/10.7554/eLife.56151 (2020).

63. Zhang, J., Huang, Z., Tumati, S. & Northoff, G. Rest-task modulation of fMRI-derived global signal topography is mediated by transient coactivation patterns. PLOS Biology 18, e3000733. https://doi.org/10.1371/journal.pbio.3000733 (July 2020).

64. Alboukadel Kassambara. rstatix: Pipe-Friendly Framework for Basic Statistical Tests. R package version 0.7.0 R package (2021).

65. Cieri, F., Zhuang, X., Caldwell, J. Z. K. & Cordes, D. Brain Entropy During Aging Through a Free Energy Principle Approach 2021. https://www.frontiersin.org/article/10.3389/fnhum.2021.647513.

66. Damiani, S. et al. From local to global and back: An exploratory study on cross-scale desynchronization in schizophrenia and its relation to thought disorders. Schizophrenia Research 231, 10–12. ISSN: 0920-9964. https://www.sciencedirect.com/science/article/pii/S0920996421001055 (2021).

67. Golesorkhi, M., Gomez-Pilar, J., Tumati, S., Fraser, M. & Northoff, G. Temporal hierarchy of intrinsic neural timescales converges with spatial core-periphery organization. Communications Biology 4, 277. ISSN: 2399-3642. https://doi.org/10.1038/s42003-021-01785-z (2021).

68. Ibaceta, M. & Madrid, H. P. Personality and Mind-Wandering Self-Perception: The Role of Meta-Awareness. Frontiers in Psychology 12. ISSN: 1664-1078. https://www.frontiersin.org/article/10.3389/fpsyg.2021.581129 (2021).

69. Omidvarnia, A. et al. Temporal complexity of fMRI is reproducible and correlates with higher order cognition. NeuroImage 230, 117760. ISSN: 1053-8119. https://www.sciencedirect.com/science/article/pii/S1053811921000379 (2021).

70. Scalabrini, A., Wolman, A. & Northoff, G. The Self and Its Right Insula—Differential Topography and Dynamic of Right vs. Left Insula 2021.

71. Smallwood, J. et al. The neural correlates of ongoing conscious thought. eng. iScience 24, 102132. ISSN: 2589-0042 (Electronic) (Mar. 2021).

72. Çatal, Y., Gomez-Pilar, J. & Northoff, G. Intrinsic Dynamics and Topography of Sensory Input Systems. Cerebral Cortex, bhab504. ISSN: 1047-3211. https://doi.org/10.1093/cercor/bhab504 (Jan. 2022).

73. David, R., Hayes, A. & Simon, C. broom: Convert Statistical Objects into Tidy Tibbles. R package version 0.8.0 (2022).

74. Golesorkhi, M. et al. From temporal to spatial topography: hierarchy of neural dynamics in higher- and lower-order networks shapes their complexity. Cerebral Cortex, bhac042. ISSN: 1047-3211. https://doi.org/10.1093/cercor/bhac042 (Feb. 2022).

75. Iannone, R., Cheng, J. & Schloerke, B. gt: Easily Create Presentation-Ready Display Tables. R package version 0.6.0 (2022).

76. Müller, K. & Wickham, H. tibble: Simple Data Frames. R package version 3.1.7 (2022).

77. R Core Team. R: A Language and Environment for Statistical Computing. R Foundation for Statistical Computing (2022).

78. Rostami, S. et al. Slow and Powerless Thought Dynamic Relates to Brooding in Unipolar and Bipolar Depression. eng. Psychopathology, 1–15. ISSN: 1423-033X (Electronic) (May 2022).

79. Scalabrini, A. et al. The self and its internal thought: In search for a psychological baseline. eng. Consciousness and cognition 97, 103244. ISSN: 1090-2376 (Electronic) (Jan. 2022).

80. Sleegers, W. tidystats: Save Output of Statistical Tests. R package version 0.5.1 (2022).

81. Smith, D., Wolff, A., Wolman, A., Ignaszewski, J. & Northoff, G. Temporal continuity of self: Long autocorrelation windows mediate self-specificity. eng. NeuroImage 257, 119305. ISSN: 1095-9572 (Electronic) (May 2022).

82. Wainio-Theberge, S., Wolff, A., Gomez-Pilar, J., Zhang, J. & Northoff, G. Variability and task-responsiveness of electrophysiological dynamics: Scalefree stability and oscillatory flexibility. eng. NeuroImage 256, 119245. ISSN: 1095-9572 (Electronic) (Aug. 2022).

83. Wolff, A. et al. Intrinsic neural timescales: temporal integration and segregation. eng. Trends in cognitive sciences 26, 159–173. ISSN: 1879-307X (Electronic) (Feb. 2022).

