## Supplementary Material for "The Intrinsic Hierarchy of Self – Converging Topography and Dynamics"

[1]The Royal's Institute of Mental Health Research & University of Ottawa. Brain and Mind Research Institute, Centre for Neural Dynamics, Faculty of Medicine, University of Ottawa, Ottawa, 145 Carling Avenue, Rm. 6435, Ottawa, Ontario K1Z 7K4, Canada
[2]Istanbul University - Cerrahpaşa, Cerrahpaşa Faculty of Medicine
[3]Shanghai Key Laboratory of Psychotic Disorders, Shanghai Mental Health Center, Shanghai Jiao Tong University School of Medicine, Shanghai, China
[4]Institute of Psychological and Behavioral Sciences, Shanghai Jiao Tong University, Shanghai, China
[5]CAS Center for Excellence in Brain Science and Intelligence Technology, Chinese Academy of Sciences, Shanghai, China
[6]Centre for Cognition and Brain Disorders, Hangzhou Normal University, Tianmu Road 305, Hangzhou, Zhejiang Province 310013, China
[7]Mental Health Centre, Zhejiang University School of Medicine, Tianmu Road 305, Hangzhou, Zhejiang Province 310013, China

June 23, 2022

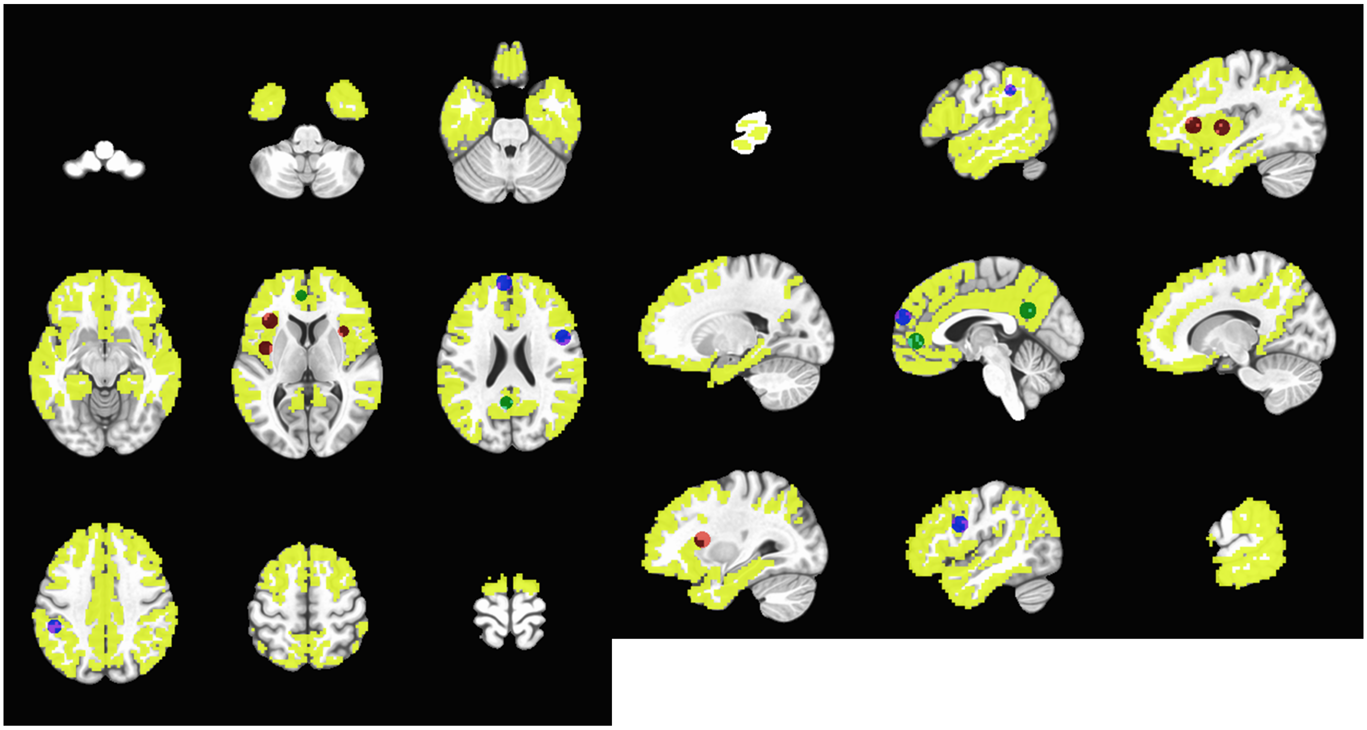

Figure 1: The ROIs we used overlaid on the transmodal (core) networks determined in [2] in atlas [1]
.

### 1 Comparison of Three Layers in Rest

SHANGHAI REST KRUSKAL-WALLIS TESTS

| | Chi-Squared | p | eta-squared |
|---|---|---|---|
| PLE | 36.51911 | *** | 0.3027992 |
| MF | 44.86776 | *** | 0.3760330 |
| SE | 30.06122 | *** | 0.2461511 |
| ACW-0 | 24.97297 | *** | 0.2015173 |
| LZC | 18.67736 | *** | 0.1462926 |

Table S1. Chi-Squared and Eta-Squared values for Kruskal-Wallis tests in the Shanghai dataset resting state among three self layers

#### SHANGHAI REST PAIRWISE COMPARISONS

| Measure | group1 | group2 | effsize | n1 | n2 | magnitude |
|---|---|---|---|---|---|---|
| PLE | EXTERO | INTERO | 0.3388925 | 39 | 39 | moderate |
| PLE | EXTERO | MENTAL | 0.4203625 | 39 | 39 | moderate |
| PLE | INTERO | MENTAL | 0.6421419 | 39 | 39 | large |
| MF | EXTERO | INTERO | 0.4158364 | 39 | 39 | moderate |
| MF | EXTERO | MENTAL | 0.4475192 | 39 | 39 | moderate |
| MF | INTERO | MENTAL | 0.7032444 | 39 | 39 | large |
| SE | EXTERO | INTERO | 0.2019777 | 39 | 39 | small |
| SE | EXTERO | MENTAL | 0.4328093 | 39 | 39 | moderate |
| SE | INTERO | MENTAL | 0.5900917 | 39 | 39 | large |
| ACW-0 | EXTERO | INTERO | 0.3128256 | 39 | 39 | moderate |
| ACW-0 | EXTERO | MENTAL | 0.2810030 | 39 | 39 | small |
| ACW-0 | INTERO | MENTAL | 0.5538638 | 39 | 39 | large |
| LZC | EXTERO | INTERO | 0.2574850 | 39 | 39 | small |
| LZC | EXTERO | MENTAL | 0.3080445 | 39 | 39 | moderate |
| LZC | INTERO | MENTAL | 0.4475863 | 39 | 39 | moderate |

Table S2. Effect estimates for pairwise Wilcoxon tests in the Shanghai dataset resting state among three self layers

#### UCLA REST KRUSKAL-WALLIS TESTS

| | Chi-Squared | p | eta-squared |
|---|---|---|---|
| PLE | 83.16268 | *** | 0.2505021 |
| MF | 115.04683 | *** | 0.3489100 |
| SE | 106.68267 | *** | 0.3230947 |
| ACW-0 | 41.40380 | *** | 0.1216167 |
| LZC | 44.53531 | *** | 0.1312818 |

Table S3. Chi-Squared and Eta-Squared values for Kruskal-Wallis tests in the UCLA dataset resting state among three self layers

#### UCLA REST PAIRWISE COMPARISONS

| Measure | group1 | group2 | effsize | n1 | n2 | magnitude |
|---|---|---|---|---|---|---|
| PLE | EXTERO | INTERO | 0.16210406 | 109 | 109 | small |
| PLE | EXTERO | MENTAL | 0.48151233 | 109 | 109 | moderate |
| PLE | INTERO | MENTAL | 0.56165530 | 109 | 109 | large |
| MF | EXTERO | INTERO | 0.15526791 | 109 | 109 | small |
| MF | EXTERO | MENTAL | 0.58696360 | 109 | 109 | large |
| MF | INTERO | MENTAL | 0.65154342 | 109 | 109 | large |
| SE | EXTERO | INTERO | 0.08443374 | 109 | 109 | small |
| SE | EXTERO | MENTAL | 0.59859961 | 109 | 109 | large |
| SE | INTERO | MENTAL | 0.60776296 | 109 | 109 | large |

| | | | | | | |
|---|---|---|---|---|---|---|
| ACW-0 | EXTERO | INTERO | 0.23506585 | 109 | 109 | small |
| ACW-0 | EXTERO | MENTAL | 0.22091112 | 109 | 109 | small |
| ACW-0 | INTERO | MENTAL | 0.42743397 | 109 | 109 | moderate |
| LZC | EXTERO | INTERO | 0.10243740 | 109 | 109 | small |
| LZC | EXTERO | MENTAL | 0.36570355 | 109 | 109 | moderate |
| LZC | INTERO | MENTAL | 0.40290502 | 109 | 109 | moderate |

Table S4. Effect estimates for pairwise Wilcoxon tests in UCLA dataset resting state among three self layers

### 2 Comparison of Three Layers in Task

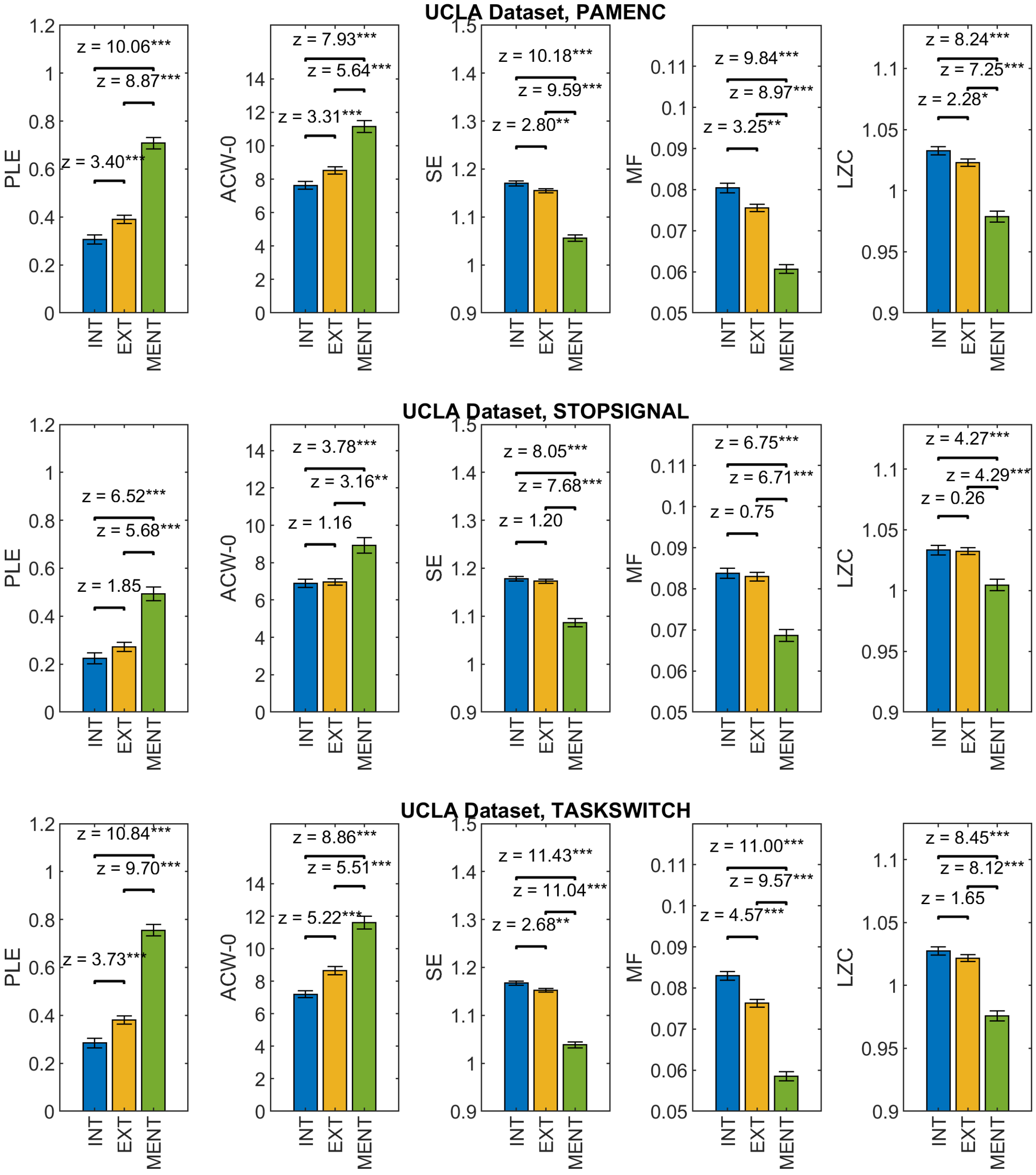


Figure 2: Pairwise comparison of the three self layers using Wilcoxon tests. Asterixes denote significance.

#### SHANGHAI TASK

| | Chi-Squared | p | eta-squared |
|---|---|---|---|
| PLE | 37.98158 | *** | 0.2855681 |
| MF | 41.54838 | *** | 0.3138760 |
| SE | 23.26676 | *** | 0.1687838 |
| ACW-0 | 21.51917 | *** | 0.1549140 |
| LZC | 28.18659 | *** | 0.2078301 |

Table S5. Chi-Squared and Eta-Squared values for Kruskal-Wallis tests in the Shanghai dataset task state among three self layers

| Measure | group1 | group2 | effsize | n1 | n2 | magnitude |
|---|---|---|---|---|---|---|
| PLE | EXTERO | INTERO | 0.1215413 | 43 | 43 | small |
| PLE | EXTERO | MENTAL | 0.5313357 | 43 | 43 | large |
| PLE | INTERO | MENTAL | 0.6030497 | 43 | 43 | large |
| MF | EXTERO | INTERO | 0.2360974 | 43 | 43 | small |
| MF | EXTERO | MENTAL | 0.5229535 | 43 | 43 | large |
| MF | INTERO | MENTAL | 0.6309902 | 43 | 43 | large |
| SE | EXTERO | INTERO | 0.1466878 | 43 | 43 | small |
| SE | EXTERO | MENTAL | 0.3953584 | 43 | 43 | moderate |
| SE | INTERO | MENTAL | 0.4773173 | 43 | 43 | moderate |
| ACW-0 | EXTERO | INTERO | 0.1923881 | 43 | 43 | small |
| ACW-0 | EXTERO | MENTAL | 0.3585141 | 43 | 43 | moderate |
| ACW-0 | INTERO | MENTAL | 0.4598354 | 43 | 43 | moderate |
| LZC | EXTERO | INTERO | 0.1859256 | 43 | 43 | small |
| LZC | EXTERO | MENTAL | 0.4280991 | 43 | 43 | moderate |
| LZC | INTERO | MENTAL | 0.5240085 | 43 | 43 | large |

Table S6. Effect estimates for Pairwise Wilcoxon tests in the Shanghai dataset task state among three self layers

#### UCLA TASKS

| | Chi-Squared | p | eta-squared |
|---|---|---|---|
| Bart PLE | 126.79919 | *** | 0.38164888 |
| Bart MF | 124.26874 | *** | 0.37391051 |
| Bart SE | 134.72077 | *** | 0.40587391 |
| Bart ACW-0 | 70.47776 | *** | 0.20941211 |
| Bart LZC | 83.63655 | *** | 0.24965305 |
| Pamenc PLE | 85.39286 | *** | 0.36100807 |
| Pamenc MF | 93.28214 | *** | 0.39516078 |
| Pamenc SE | 105.96873 | *** | 0.45008109 |
| Pamenc ACW-0 | 39.59055 | *** | 0.16272965 |
| Pamenc LZC | 60.02045 | *** | 0.25117079 |

| | | | |
|---|---|---|---|
| Pamret PLE | 51.98484 | *** | 0.21638460 |
| Pamret MF | 60.86435 | *** | 0.25482401 |
| Pamret SE | 83.58926 | *** | 0.35320027 |
| Pamret ACW-0 | 17.66097 | *** | 0.06779643 |
| Pamret LZC | 24.60406 | *** | 0.09785309 |
| Scap PLE | 149.08600 | *** | 0.45396913 |
| Scap MF | 153.77217 | *** | 0.46843262 |
| Scap SE | 172.25503 | *** | 0.52547848 |
| Scap ACW-0 | 88.87596 | *** | 0.26813569 |
| Scap LZC | 93.30068 | *** | 0.28179221 |
| Stopsignal PLE | 125.63460 | *** | 0.38515450 |
| Stopsignal MF | 138.50348 | *** | 0.42524450 |
| Stopsignal SE | 134.83770 | *** | 0.41382460 |
| Stopsignal ACW-0 | 75.85159 | *** | 0.23006726 |
| Stopsignal LZC | 95.87437 | *** | 0.29244353 |
| Taskswitch PLE | 96.97362 | *** | 0.29865918 |
| Taskswitch MF | 108.46619 | *** | 0.33479936 |
| Taskswitch SE | 108.89088 | *** | 0.33613485 |
| Taskswitch ACW-0 | 35.60019 | *** | 0.10566096 |
| Taskswitch LZC | 44.77670 | *** | 0.13451791 |

Table S7. Chi-Squared and Eta-Squared values for Kruskal-Wallis tests in the UCLA dataset Task States among three self layers.

UCLA TASKS PAIRWISE COMPARISONS

| Measure | Layer1 | Layer2 | effsize | n1 | n2 | magnitude |
|---|---|---|---|---|---|---|
| Bart PLE | EXTERO | INTERO | 0.22951250 | 110 | 110 | small |
| Bart PLE | EXTERO | MENTAL | 0.59813213 | 110 | 110 | large |
| Bart PLE | INTERO | MENTAL | 0.67839724 | 110 | 110 | large |
| Bart MF | EXTERO | INTERO | 0.21937224 | 110 | 110 | small |
| Bart MF | EXTERO | MENTAL | 0.60498751 | 110 | 110 | large |
| Bart MF | INTERO | MENTAL | 0.66368673 | 110 | 110 | large |
| Bart SE | EXTERO | INTERO | 0.18866584 | 110 | 110 | small |
| Bart SE | EXTERO | MENTAL | 0.64654827 | 110 | 110 | large |
| Bart SE | INTERO | MENTAL | 0.68625236 | 110 | 110 | large |
| Bart ACW-0 | EXTERO | INTERO | 0.22326086 | 110 | 110 | small |
| Bart ACW-0 | EXTERO | MENTAL | 0.38060273 | 110 | 110 | moderate |
| Bart ACW-0 | INTERO | MENTAL | 0.53484585 | 110 | 110 | large |
| Bart LZC | EXTERO | INTERO | 0.15349086 | 110 | 110 | small |
| Bart LZC | EXTERO | MENTAL | 0.48904920 | 110 | 110 | moderate |
| Bart LZC | INTERO | MENTAL | 0.55556549 | 110 | 110 | large |
| Pamenc PLE | EXTERO | INTERO | 0.24433577 | 78 | 78 | small |
| Pamenc PLE | EXTERO | MENTAL | 0.56500873 | 78 | 78 | large |
| Pamenc PLE | INTERO | MENTAL | 0.66858893 | 78 | 78 | large |

| | | | | | | |
|---|---|---|---|---|---|---|
| Pamenc MF | EXTERO | INTERO | 0.23894392 | 78 | 78 | small |
| Pamenc MF | EXTERO | MENTAL | 0.60814350 | 78 | 78 | large |
| Pamenc MF | INTERO | MENTAL | 0.69015631 | 78 | 78 | large |
| Pamenc SE | EXTERO | INTERO | 0.04795905 | 78 | 78 | small |
| Pamenc SE | EXTERO | MENTAL | 0.71739933 | 78 | 78 | large |
| Pamenc SE | INTERO | MENTAL | 0.70775076 | 78 | 78 | large |
| Pamenc ACW-0 | EXTERO | INTERO | 0.20391318 | 78 | 78 | small |
| Pamenc ACW-0 | EXTERO | MENTAL | 0.32709873 | 78 | 78 | moderate |
| Pamenc ACW-0 | INTERO | MENTAL | 0.48070022 | 78 | 78 | moderate |
| Pamenc LZC | EXTERO | INTERO | 0.05230058 | 78 | 78 | small |
| Pamenc LZC | EXTERO | MENTAL | 0.56218081 | 78 | 78 | large |
| Pamenc LZC | INTERO | MENTAL | 0.50650684 | 78 | 78 | large |
| Pamret PLE | EXTERO | INTERO | 0.14813388 | 78 | 78 | small |
| Pamret PLE | EXTERO | MENTAL | 0.45461777 | 78 | 78 | moderate |
| Pamret PLE | INTERO | MENTAL | 0.52187396 | 78 | 78 | large |
| Pamret MF | EXTERO | INTERO | 0.06016165 | 78 | 78 | small |
| Pamret MF | EXTERO | MENTAL | 0.53748193 | 78 | 78 | large |
| Pamret MF | INTERO | MENTAL | 0.54088731 | 78 | 78 | large |
| Pamret SE | EXTERO | INTERO | 0.09620189 | 78 | 78 | small |
| Pamret SE | EXTERO | MENTAL | 0.61495425 | 78 | 78 | large |
| Pamret SE | INTERO | MENTAL | 0.64503507 | 78 | 78 | large |
| Pamret ACW-0 | EXTERO | INTERO | 0.09307332 | 78 | 78 | small |
| Pamret ACW-0 | EXTERO | MENTAL | 0.25334053 | 78 | 78 | small |
| Pamret ACW-0 | INTERO | MENTAL | 0.30313783 | 78 | 78 | moderate |
| Pamret LZC | EXTERO | INTERO | 0.02127546 | 78 | 78 | small |
| Pamret LZC | EXTERO | MENTAL | 0.34401682 | 78 | 78 | moderate |
| Pamret LZC | INTERO | MENTAL | 0.34217199 | 78 | 78 | moderate |
| Scap PLE | EXTERO | INTERO | 0.25271943 | 109 | 109 | small |
| Scap PLE | EXTERO | MENTAL | 0.65721597 | 109 | 109 | large |
| Scap PLE | INTERO | MENTAL | 0.73401359 | 109 | 109 | large |
| Scap MF | EXTERO | INTERO | 0.30973584 | 109 | 109 | moderate |
| Scap MF | EXTERO | MENTAL | 0.64848897 | 109 | 109 | large |
| Scap MF | INTERO | MENTAL | 0.74506779 | 109 | 109 | large |
| Scap SE | EXTERO | INTERO | 0.18130346 | 109 | 109 | small |
| Scap SE | EXTERO | MENTAL | 0.74783134 | 109 | 109 | large |
| Scap SE | INTERO | MENTAL | 0.77415779 | 109 | 109 | large |
| Scap ACW-0 | EXTERO | INTERO | 0.35339589 | 109 | 109 | moderate |
| Scap ACW-0 | EXTERO | MENTAL | 0.37355918 | 109 | 109 | moderate |
| Scap ACW-0 | INTERO | MENTAL | 0.60017740 | 109 | 109 | large |
| Scap LZC | EXTERO | INTERO | 0.11183292 | 109 | 109 | small |
| Scap LZC | EXTERO | MENTAL | 0.54969480 | 109 | 109 | large |
| Scap LZC | INTERO | MENTAL | 0.57242119 | 109 | 109 | large |
| Stopsignal PLE | EXTERO | INTERO | 0.23319265 | 108 | 108 | small |
| Stopsignal PLE | EXTERO | MENTAL | 0.59957474 | 108 | 108 | large |
| Stopsignal PLE | INTERO | MENTAL | 0.68179960 | 108 | 108 | large |
| Stopsignal MF | EXTERO | INTERO | 0.26015647 | 108 | 108 | small |

| | | | | | | |
|---|---|---|---|---|---|---|
| Stopsignal MF | EXTERO | MENTAL | 0.64772443 | 108 | 108 | large |
| Stopsignal MF | INTERO | MENTAL | 0.69839272 | 108 | 108 | large |
| Stopsignal SE | EXTERO | INTERO | 0.15422716 | 108 | 108 | small |
| Stopsignal SE | EXTERO | MENTAL | 0.65587284 | 108 | 108 | large |
| Stopsignal SE | INTERO | MENTAL | 0.69661488 | 108 | 108 | large |
| Stopsignal ACW-0 | EXTERO | INTERO | 0.26544789 | 108 | 108 | small |
| Stopsignal ACW-0 | EXTERO | MENTAL | 0.37715159 | 108 | 108 | moderate |
| Stopsignal ACW-0 | INTERO | MENTAL | 0.56200013 | 108 | 108 | large |
| Stopsignal LZC | EXTERO | INTERO | 0.07238908 | 108 | 108 | small |
| Stopsignal LZC | EXTERO | MENTAL | 0.55122683 | 108 | 108 | large |
| Stopsignal LZC | INTERO | MENTAL | 0.59345025 | 108 | 108 | large |
| Taskswitch PLE | EXTERO | INTERO | 0.18209921 | 107 | 107 | small |
| Taskswitch PLE | EXTERO | MENTAL | 0.53844917 | 107 | 107 | large |
| Taskswitch PLE | INTERO | MENTAL | 0.60033120 | 107 | 107 | large |
| Taskswitch MF | EXTERO | INTERO | 0.16489298 | 107 | 107 | small |
| Taskswitch MF | EXTERO | MENTAL | 0.57391814 | 107 | 107 | large |
| Taskswitch MF | INTERO | MENTAL | 0.63715856 | 107 | 107 | large |
| Taskswitch SE | EXTERO | INTERO | 0.02030033 | 107 | 107 | small |
| Taskswitch SE | EXTERO | MENTAL | 0.62463122 | 107 | 107 | large |
| Taskswitch SE | INTERO | MENTAL | 0.60983991 | 107 | 107 | large |
| Taskswitch ACW-0 | EXTERO | INTERO | 0.19674271 | 107 | 107 | small |
| Taskswitch ACW-0 | EXTERO | MENTAL | 0.23571333 | 107 | 107 | small |
| Taskswitch ACW-0 | INTERO | MENTAL | 0.39409352 | 107 | 107 | moderate |
| Taskswitch LZC | EXTERO | INTERO | 0.05482110 | 107 | 107 | small |
| Taskswitch LZC | EXTERO | MENTAL | 0.42440026 | 107 | 107 | moderate |
| Taskswitch LZC | INTERO | MENTAL | 0.36063689 | 107 | 107 | moderate |

Table S8. Effect estimates for pairwise Wilcoxon tests in the UCLA dataset task states among three self layers

### 3 Rest-Task Differences

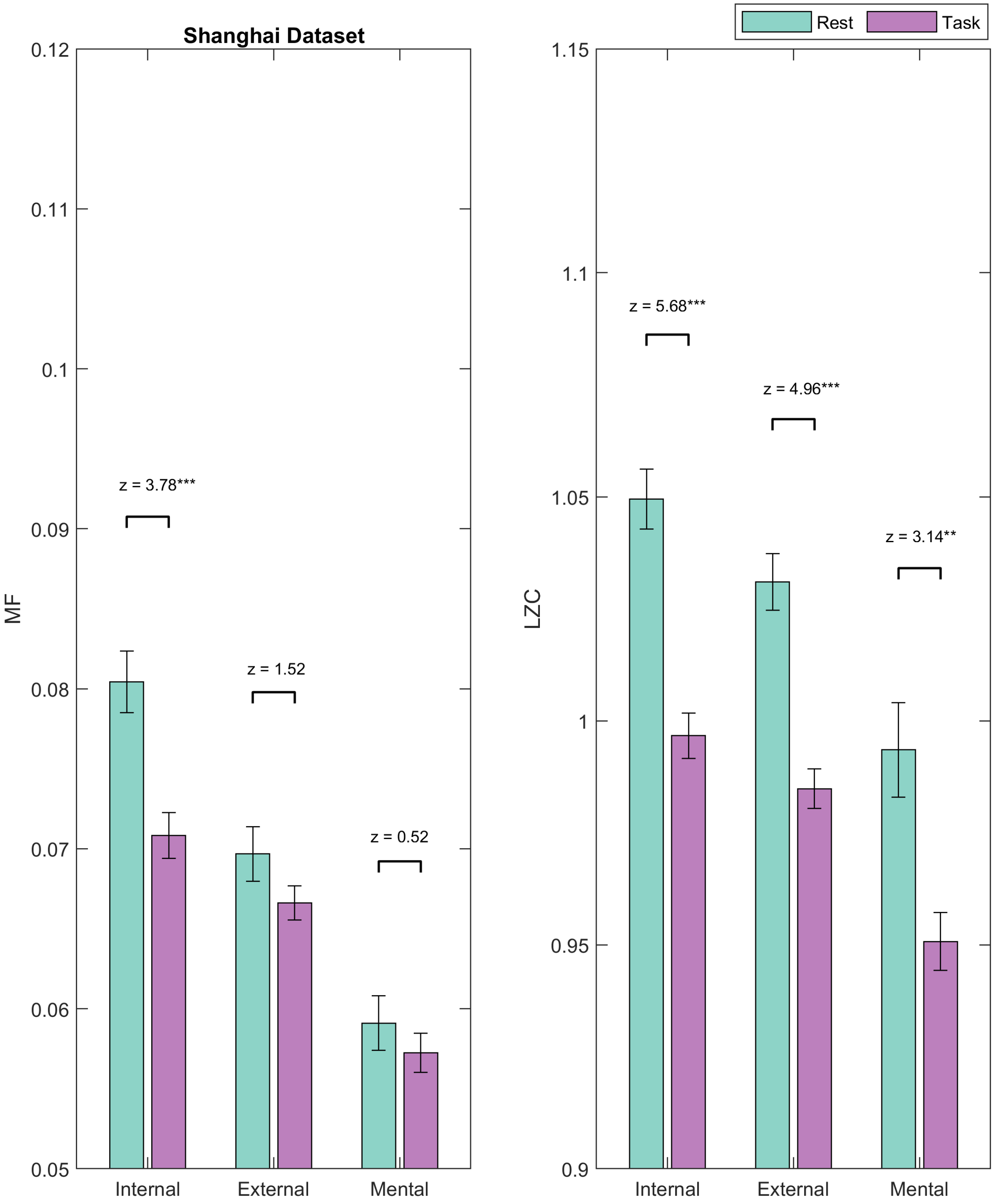


Figure 3: Rest-Task comparison of dynamic measures in the Shanghai dataset

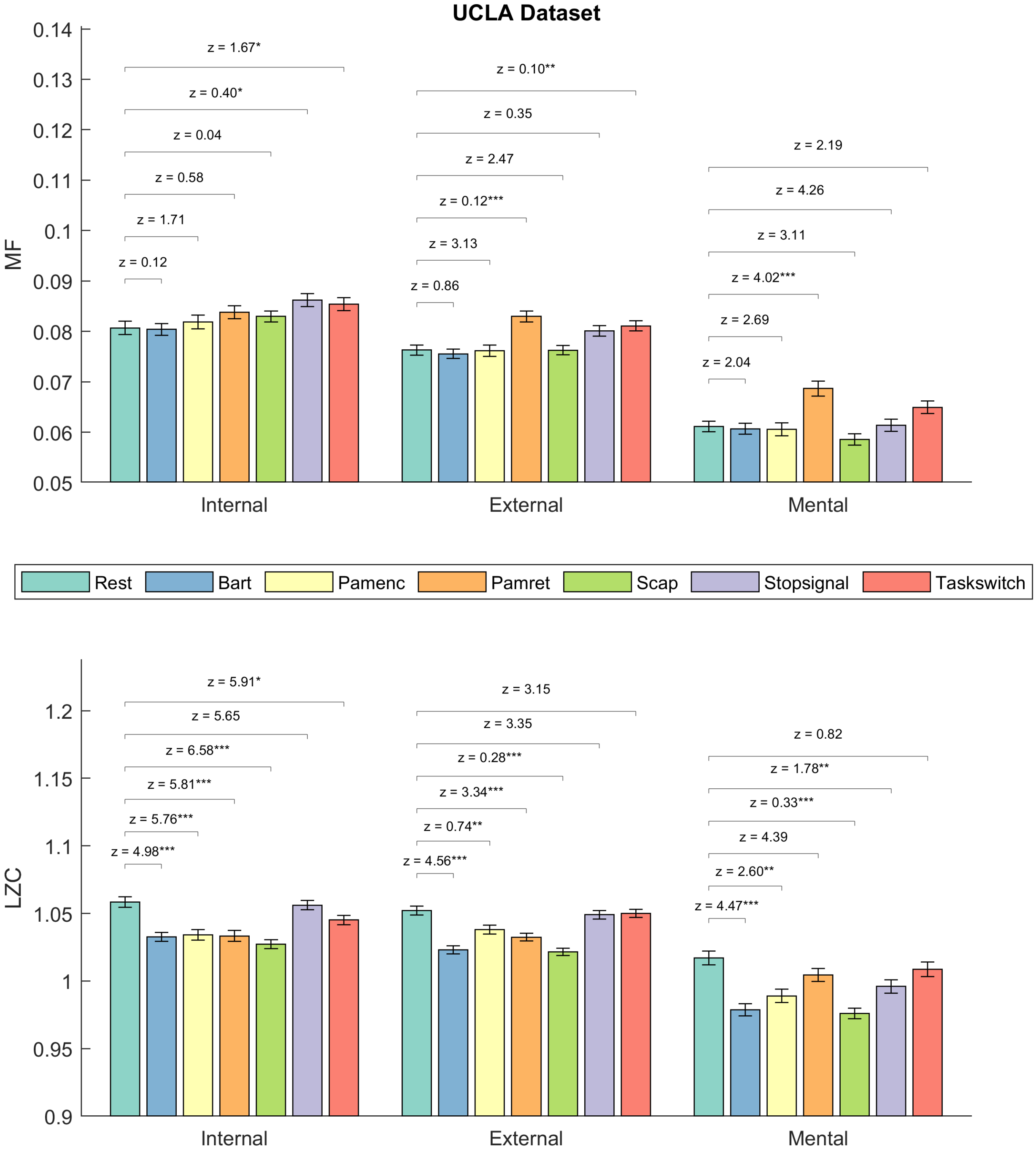


Figure 4: Rest-Task comparison of dynamic measures in the UCLA dataset

SHANGHAI REST - TASK COMPARSIONS

| group1 | group2 | effsize | n1 | n2 | magnitude |
|---|---|---|---|---|---|
| INTERO PLE Rest | Task | 0.55318395 | 39 | 43 | large |
| INTERO MF Rest | Task | 0.41783588 | 39 | 43 | moderate |
| INTERO SE Rest | Task | 0.44962217 | 39 | 43 | moderate |
| INTERO ACW-0 Rest | Task | 0.36842249 | 39 | 43 | moderate |
| INTERO LZC Rest | Task | 0.62772451 | 39 | 43 | large |
| EXTERO PLE Rest | Task | 0.28351318 | 39 | 43 | small |
| EXTERO MF Rest | Task | 0.16867240 | 39 | 43 | small |
| EXTERO SE Rest | Task | 0.39425251 | 39 | 43 | moderate |
| EXTERO ACW-0 Rest | Task | 0.09954367 | 39 | 43 | small |
| EXTERO LZC Rest | Task | 0.54779543 | 39 | 43 | large |
| MENTAL PLE Rest | Task | 0.20250942 | 39 | 43 | small |
| MENTAL MF Rest | Task | 0.05793307 | 39 | 43 | small |
| MENTAL SE Rest | Task | 0.39117641 | 39 | 43 | moderate |
| MENTAL ACW-0 Rest | Task | 0.06876315 | 39 | 43 | small |
| MENTAL LZC Rest | Task | 0.34738646 | 39 | 43 | moderate |

Table S9. Rest-Task differences of three self layers in the Shanghai dataset

UCLA REST - TASK COMPARISONS

| group1 | group2 | effsize | n1 | n2 | magnitude |
|---|---|---|---|---|---|
| INTERO PLE Rest | Bart | 0.083161717 | 110 | 109 | small |
| EXTERO PLE Rest | Bart | 0.133750560 | 110 | 109 | small |
| MENTAL PLE Rest | Bart | 0.142686481 | 110 | 109 | small |
| INTERO MF Rest | Bart | 0.008071154 | 110 | 109 | small |
| EXTERO MF Rest | Bart | 0.039491006 | 110 | 109 | small |
| MENTAL MF Rest | Bart | 0.027240146 | 110 | 109 | small |
| INTERO SE Rest | Bart | 0.029113807 | 110 | 109 | small |
| EXTERO SE Rest | Bart | 0.123949873 | 110 | 109 | small |
| MENTAL SE Rest | Bart | 0.188374981 | 110 | 109 | small |
| INTERO ACW-0 Rest | Bart | 0.062386189 | 110 | 109 | small |
| EXTERO ACW-0 Rest | Bart | 0.076471373 | 110 | 109 | small |
| MENTAL ACW-0 Rest | Bart | 0.062572007 | 110 | 109 | small |
| INTERO LZC Rest | Bart | 0.336751752 | 110 | 109 | moderate |
| EXTERO LZC Rest | Bart | 0.392576428 | 110 | 109 | moderate |
| MENTAL LZC Rest | Bart | 0.382043577 | 110 | 109 | moderate |
| INTERO PLE Rest | Pamenc | 0.014626931 | 109 | 78 | small |
| EXTERO PLE Rest | Pamenc | 0.080949041 | 109 | 78 | small |
| MENTAL PLE Rest | Pamenc | 0.089164167 | 109 | 78 | small |
| INTERO MF Rest | Pamenc | 0.062715470 | 109 | 78 | small |
| EXTERO MF Rest | Pamenc | 0.008615863 | 109 | 78 | small |
| MENTAL MF Rest | Pamenc | 0.025446852 | 109 | 78 | small |
| INTERO SE Rest | Pamenc | 0.065520635 | 109 | 78 | small |

| | | | | | |
|---|---|---|---|---|---|
| EXTERO SE Rest | Pamenc | 0.028252017 | 109 | 78 | small |
| MENTAL SE Rest | Pamenc | 0.186343090 | 109 | 78 | small |
| INTERO ACW-0 Rest | Pamenc | 0.113712363 | 109 | 78 | small |
| EXTERO ACW-0 Rest | Pamenc | 0.146656969 | 109 | 78 | small |
| MENTAL ACW-0 Rest | Pamenc | 0.002745653 | 109 | 78 | small |
| INTERO LZC Rest | Pamenc | 0.333910435 | 109 | 78 | moderate |
| EXTERO LZC Rest | Pamenc | 0.244431703 | 109 | 78 | small |
| MENTAL LZC Rest | Pamenc | 0.244933318 | 109 | 78 | small |
| INTERO PLE Rest | Pamret | 0.073936129 | 109 | 78 | small |
| EXTERO PLE Rest | Pamret | 0.153081850 | 109 | 78 | small |
| MENTAL PLE Rest | Pamret | 0.245652289 | 109 | 78 | small |
| INTERO MF Rest | Pamret | 0.149475210 | 109 | 78 | small |
| EXTERO MF Rest | Pamret | 0.293941197 | 109 | 78 | small |
| MENTAL MF Rest | Pamret | 0.311373293 | 109 | 78 | moderate |
| INTERO SE Rest | Pamret | 0.030656444 | 109 | 78 | small |
| EXTERO SE Rest | Pamret | 0.072934285 | 109 | 78 | small |
| MENTAL SE Rest | Pamret | 0.047487433 | 109 | 78 | small |
| INTERO ACW-0 Rest | Pamret | 0.238902297 | 109 | 78 | small |
| EXTERO ACW-0 Rest | Pamret | 0.409756003 | 109 | 78 | moderate |
| MENTAL ACW-0 Rest | Pamret | 0.283545942 | 109 | 78 | small |
| INTERO LZC Rest | Pamret | 0.326920777 | 109 | 78 | moderate |
| EXTERO LZC Rest | Pamret | 0.321171489 | 109 | 78 | moderate |
| MENTAL LZC Rest | Pamret | 0.130396284 | 109 | 78 | small |
| INTERO PLE Rest | Scap | 0.046471285 | 109 | 109 | small |
| EXTERO PLE Rest | Scap | 0.103633148 | 109 | 109 | small |
| MENTAL PLE Rest | Scap | 0.238174427 | 109 | 109 | small |
| INTERO MF Rest | Scap | 0.115705500 | 109 | 109 | small |
| EXTERO MF Rest | Scap | 0.002981726 | 109 | 109 | small |
| MENTAL MF Rest | Scap | 0.113232850 | 109 | 109 | small |
| INTERO SE Rest | Scap | 0.069597840 | 109 | 109 | small |
| EXTERO SE Rest | Scap | 0.148140857 | 109 | 109 | small |
| MENTAL SE Rest | Scap | 0.348716452 | 109 | 109 | moderate |
| INTERO ACW-0 Rest | Scap | 0.160320311 | 109 | 109 | small |
| EXTERO ACW-0 Rest | Scap | 0.062622593 | 109 | 109 | small |
| MENTAL ACW-0 Rest | Scap | 0.103354072 | 109 | 109 | small |
| INTERO LZC Rest | Scap | 0.390162247 | 109 | 109 | moderate |
| EXTERO LZC Rest | Scap | 0.445565984 | 109 | 109 | moderate |
| MENTAL LZC Rest | Scap | 0.400301793 | 109 | 109 | moderate |
| INTERO PLE Rest | Stopsignal | 0.226062280 | 109 | 108 | small |
| EXTERO PLE Rest | Stopsignal | 0.184079285 | 109 | 108 | small |
| MENTAL PLE Rest | Stopsignal | 0.001467937 | 109 | 108 | small |
| INTERO MF Rest | Stopsignal | 0.212410467 | 109 | 108 | small |
| EXTERO MF Rest | Stopsignal | 0.167931979 | 109 | 108 | small |
| MENTAL MF Rest | Stopsignal | 0.006752510 | 109 | 108 | small |
| INTERO SE Rest | Stopsignal | 0.125215016 | 109 | 108 | small |
| EXTERO SE Rest | Stopsignal | 0.089397356 | 109 | 108 | small |

| | | | | | |
|---|---|---|---|---|---|
| MENTAL SE Rest | Stopsignal | 0.070901351 | 109 | 108 | small |
| INTERO ACW-0 Rest | Stopsignal | 0.196097987 | 109 | 108 | small |
| EXTERO ACW-0 Rest | Stopsignal | 0.200493711 | 109 | 108 | small |
| MENTAL ACW-0 Rest | Stopsignal | 0.040866336 | 109 | 108 | small |
| INTERO LZC Rest | Stopsignal | 0.050115006 | 109 | 108 | small |
| EXTERO LZC Rest | Stopsignal | 0.019402012 | 109 | 108 | small |
| MENTAL LZC Rest | Stopsignal | 0.213746501 | 109 | 108 | small |
| INTERO PLE Rest | Taskswitch | 0.106452414 | 109 | 107 | small |
| EXTERO PLE Rest | Taskswitch | 0.136084262 | 109 | 107 | small |
| MENTAL PLE Rest | Taskswitch | 0.105711618 | 109 | 107 | small |
| INTERO MF Rest | Taskswitch | 0.182902582 | 109 | 107 | small |
| EXTERO MF Rest | Taskswitch | 0.211497316 | 109 | 107 | small |
| MENTAL MF Rest | Taskswitch | 0.148825957 | 109 | 107 | small |
| INTERO SE Rest | Taskswitch | 0.004963335 | 109 | 107 | small |
| EXTERO SE Rest | Taskswitch | 0.068819967 | 109 | 107 | small |
| MENTAL SE Rest | Taskswitch | 0.008519156 | 109 | 107 | small |
| INTERO ACW-0 Rest | Taskswitch | 0.188543568 | 109 | 107 | small |
| EXTERO ACW-0 Rest | Taskswitch | 0.229098883 | 109 | 107 | small |
| MENTAL ACW-0 Rest | Taskswitch | 0.166791727 | 109 | 107 | small |
| INTERO LZC Rest | Taskswitch | 0.177267327 | 109 | 107 | small |
| EXTERO LZC Rest | Taskswitch | 0.022786565 | 109 | 107 | small |
| MENTAL LZC Rest | Taskswitch | 0.055758659 | 109 | 107 | small |

Table S10. Rest-Task differences of three self layers in the UCLA dataset

### 4 Comparison of Gradient Indices in Rest and Task

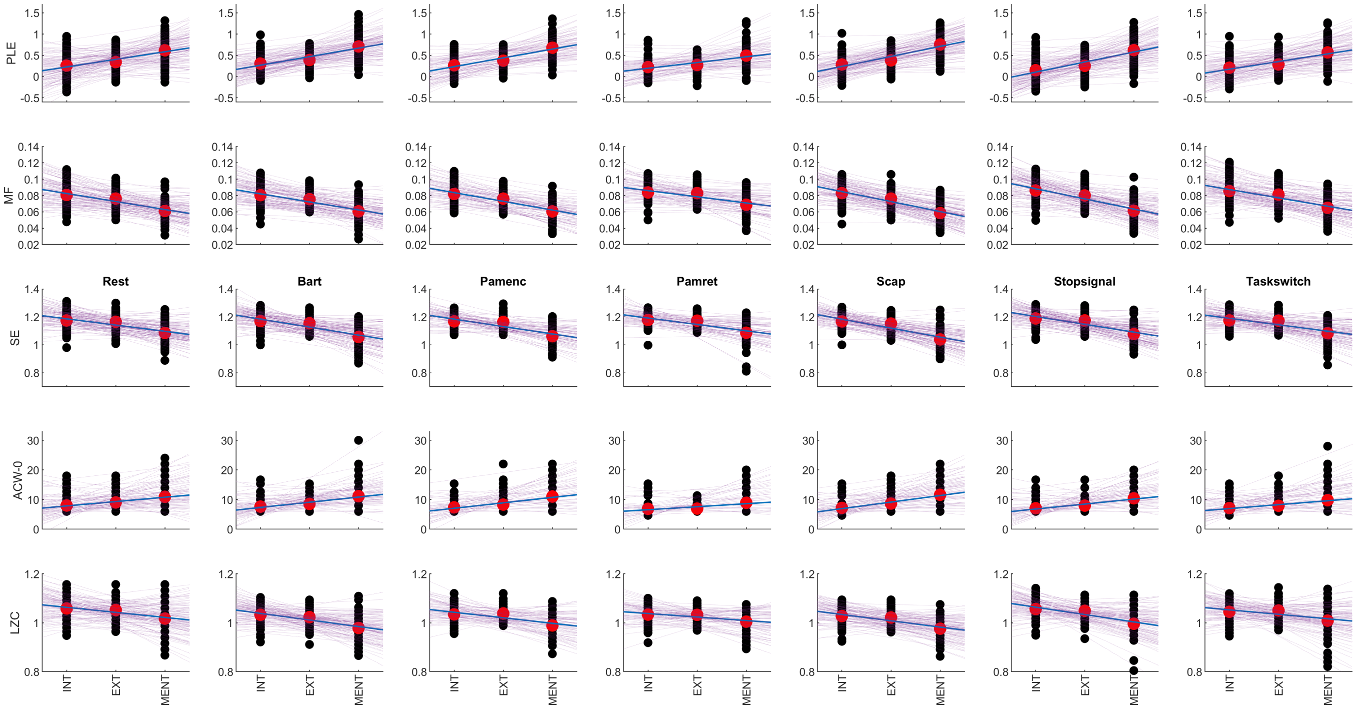


Figure 5: Demonstration of the calculation of gradient indices in the UCLA dataset

COMPARISON OF GRADIENT INDICES IN SHANGHAI

| group1 | group2 | effsize | n1 | n2 | magnitude |
|---|---|---|---|---|---|
| PLE Gradient Rest | Task | 0.28556391 | 39 | 43 | small |
| MF Gradient Rest | Task | 0.40758224 | 39 | 43 | moderate |
| SE Gradient Rest | Task | 0.13381002 | 39 | 43 | small |
| ACW-0 Gradient Rest | Task | 0.23022557 | 39 | 43 | small |
| LZC Gradient Rest | Task | 0.09177109 | 39 | 43 | small |

Table S11. Rest-Task differences of gradient indices in the Shanghai dataset

COMPARISON OF GRADIENT INDICES IN UCLA

| group1 | group2 | effsize | n1 | n2 | magnitude |
|---|---|---|---|---|---|
| Bart | PLE Gradient Rest | 0.029978574 | 110 | 109 | small |

| | | | | | |
|---|---|---|---|---|---|
| Bart | MF Gradient Rest | 0.002450172 | 110 | 109 | small |
| Bart | SE Gradient Rest | 0.131732772 | 110 | 109 | small |
| ACW-0 Gradient Rest | Bart | 0.020261510 | 109 | 110 | small |
| Bart | LZC Gradient Rest | 0.033293588 | 110 | 109 | small |
| Pamenc | PLE Gradient Rest | 0.034062715 | 78 | 109 | small |
| MF Gradient Rest | Pamenc | 0.055502189 | 109 | 78 | small |
| Pamenc | SE Gradient Rest | 0.064719159 | 78 | 109 | small |
| ACW-0 Gradient Rest | Pamenc | 0.014637047 | 109 | 78 | small |
| LZC Gradient Rest | Pamenc | 0.122025173 | 109 | 78 | small |
| Pamret | PLE Gradient Rest | 0.230824989 | 78 | 109 | small |
| MF Gradient Rest | Pamret | 0.206780719 | 109 | 78 | small |
| Pamret | SE Gradient Rest | 0.073735760 | 78 | 109 | small |
| ACW-0 Gradient Rest | Pamret | 0.271968088 | 109 | 78 | small |
| LZC Gradient Rest | Pamret | 0.220206851 | 109 | 78 | small |
| PLE Gradient Rest | Scap | 0.169667462 | 109 | 109 | small |
| MF Gradient Rest | Scap | 0.174612763 | 109 | 109 | small |
| Scap | SE Gradient Rest | 0.242974278 | 109 | 109 | small |
| ACW-0 Gradient Rest | Scap | 0.106098165 | 109 | 109 | small |
| LZC Gradient Rest | Scap | 0.064507351 | 109 | 109 | small |
| PLE Gradient Rest | Stopsignal | 0.144444989 | 109 | 108 | small |
| MF Gradient Rest | Stopsignal | 0.182170967 | 109 | 108 | small |
| SE Gradient Rest | Stopsignal | 0.133582256 | 109 | 108 | small |
| ACW-0 Gradient Rest | Stopsignal | 0.024020537 | 109 | 108 | small |
| LZC Gradient Rest | Stopsignal | 0.067525374 | 109 | 108 | small |
| PLE Gradient Rest | Taskswitch | 0.037262049 | 109 | 107 | small |
| MF Gradient Rest | Taskswitch | 0.031780157 | 109 | 107 | small |
| SE Gradient Rest | Taskswitch | 0.010297067 | 109 | 107 | small |
| ACW-0 Gradient Rest | Taskswitch | 0.108400619 | 109 | 107 | small |
| LZC Gradient Rest | Taskswitch | 0.131491913 | 109 | 107 | small |

Table S12. Rest-Task differences of gradient indices in the UCLA dataset

### 5 Network Analyses

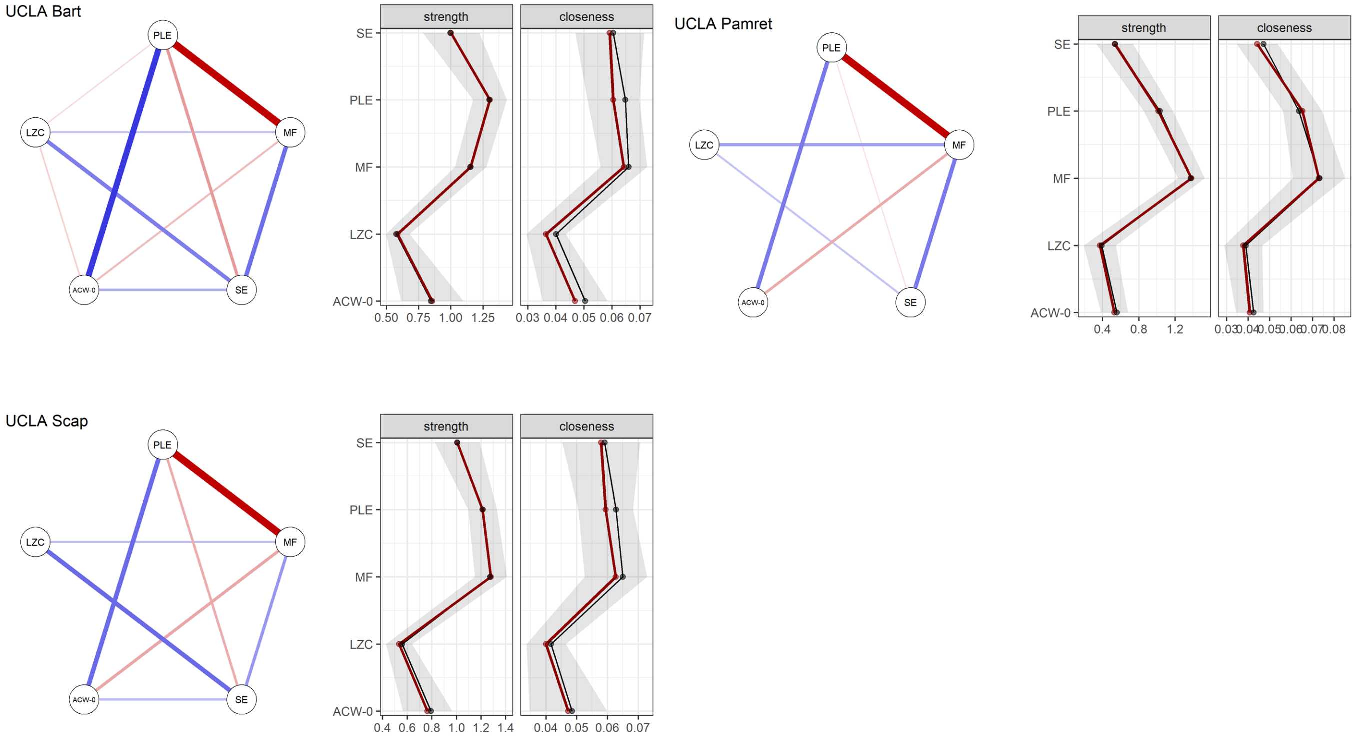


Figure 6: Additional tasks in network analysis

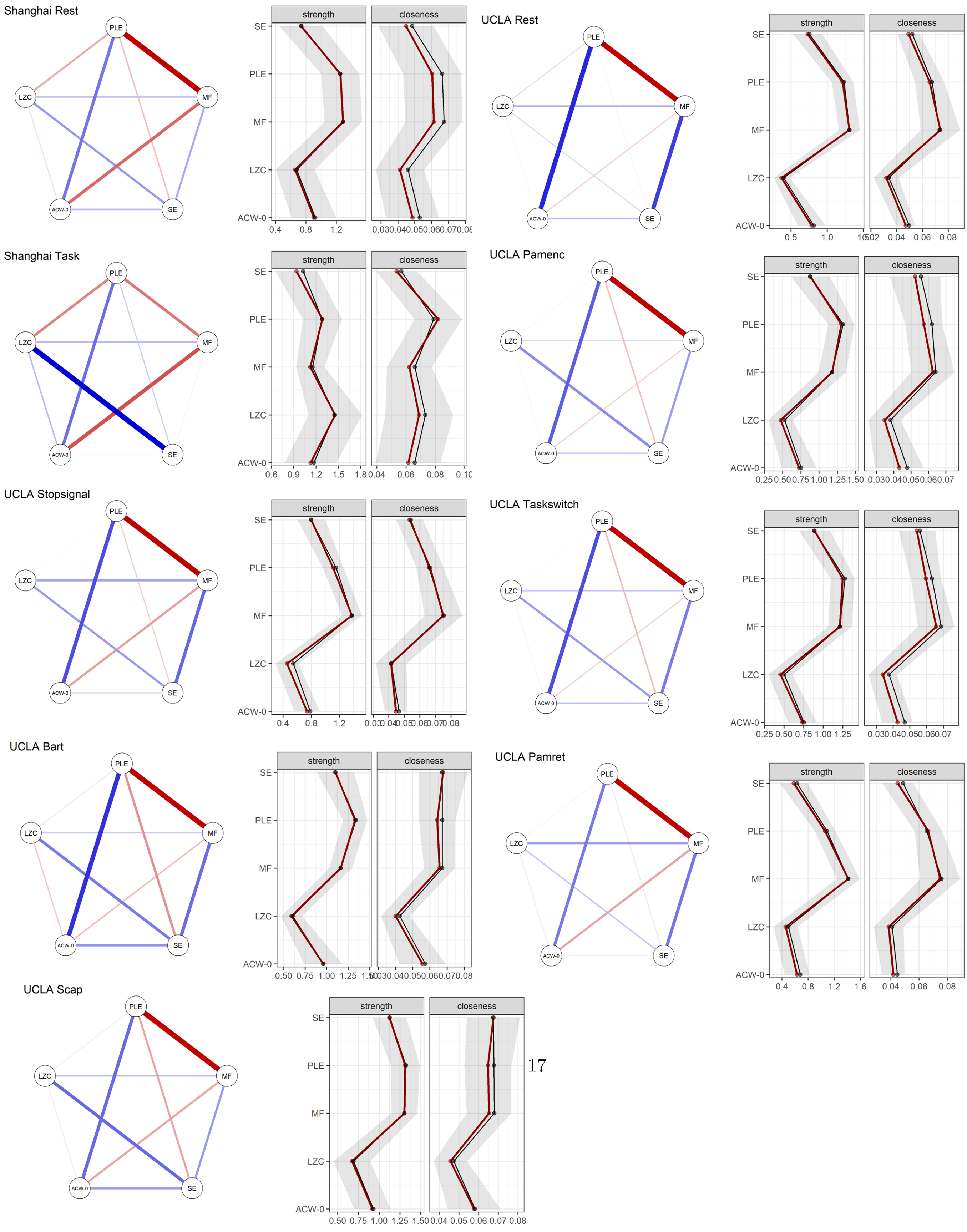
Shanghai Rest
UCLA Rest
Shanghai Task
UCLA Pamenc
UCLA Stopsignal
UCLA Taskswitch
UCLA Bart
UCLA Pamret
UCLA Scap
PLE
MF
LZC
SE
ACW-0
strength
closeness

Figure 7: Network analysis with the pcor method that doesn't depend on the sparsity assumption

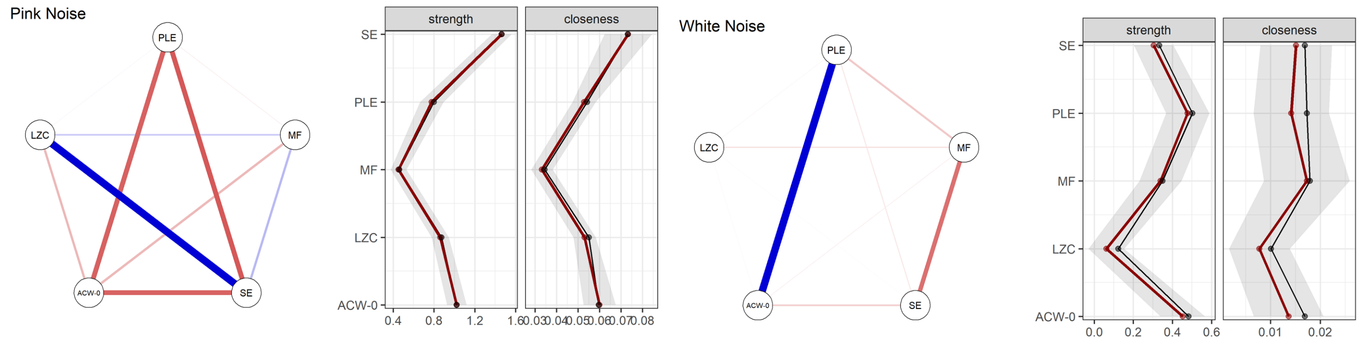


Figure 8: Network analysis of the simulated signals with the pcor method that doesn't depend on the sparsity assumption

### References


1. Ji, J. L. *et al.* Mapping the human brain's cortical-subcortical functional network organization. *NeuroImage* **185,** 35–57. ISSN: 1053-8119. `https://www.sciencedirect.com/science/article/pii/S1053811918319657` (2019).
2. Ito, T., Hearne, L. J. & Cole, M. W. A cortical hierarchy of localized and distributed processes revealed via dissociation of task activations, connectivity changes, and intrinsic timescales. *NeuroImage* **221,** 117141. ISSN: 1053-8119 (Nov. 2020).
